# CXCR4 blockade alleviates pulmonary and cardiac outcomes in early COPD

**DOI:** 10.1101/2023.03.10.529743

**Authors:** Isabelle Dupin, Pauline Henrot, Elise Maurat, Reshed Abohalaka, Sébastien Chaigne, Dounia El Hamrani, Edmée Eyraud, Renaud Prevel, Pauline Esteves, Maryline Campagnac, Marielle Dubreuil, Guillaume Cardouat, Clément Bouchet, Olga Ousova, Jean-William Dupuy, Thomas Trian, Matthieu Thumerel, Hugues Bégueret, Pierre-Olivier Girodet, Roger Marthan, Maeva Zysman, Véronique Freund-Michel, Patrick Berger

**Affiliations:** Univ. Bordeaux, Centre de Recherche Cardio-thoracique de Bordeaux, INSERM U1045, IHU Liryc, CIC 1401, Proteomics Facility, F-33600 Pessac, France; INSERM, Centre de Recherche Cardio-thoracique de Bordeaux, U1045, CIC 1401, F-33600 Pessac, France; Institut universitaire de France (IUF); CHU Bordeaux, Service d’exploration fonctionnelle respiratoire, Service de réanimation, Service de pneumologie, Service de chirurgie thoracique, Service de cardiologie-électrophysiologie et stimulation cardiaque, Service d’anatomopathologie, CIC-P 1401, F-33600 Pessac, France; Krefting Research Centre, Department of Internal Medicine and Clinical Nutrition, Institute of Medicine, Sahlgrenska Academy, University of Gothenburg - Gothenburg, Sweden

**Keywords:** COPD pathology, macrophage biology, cytokine biology

## Abstract

Chronic obstructive pulmonary disease (COPD) is a prevalent respiratory disease lacking effective treatment. Focusing on early COPD should help to discover disease modifying therapies. We aimed to examine the role of the CXCL12/CXCR4 axis in early COPD from both human samples and murine models. Blood samples and lung tissues of early COPD patients and controls were obtained in order to analyse CXCL12 and CXCR4 levels. To generate an early COPD model, ten-week-old male C57BL/6J mice were exposed to cigarette smoke (CS) for 10 weeks and intranasal instillations of polyinosinic–polycytidylic acid (poly(I:C)) for the last 5 weeks to mimic exacerbations. CXCR4 expressing cells number was increased in the blood of patients with COPD, as well as in the blood of exposed mice. Lung CXCL12 expression was higher in both early COPD patients and exposed mice. Exposed mice presented mild airway obstruction, peri-bronchial fibrosis and right heart thickening. The density of fibrocytes expressing CXCR4 was increased in the bronchial submucosa of these mice. Conditional inactivation of CXCR4 at adult stage as well as pharmacological inhibition of CXCR4 with plerixafor injections improved lung function, reduced inflammation, and protected against CS and poly-(I:C)-induced airway and cardiac remodeling. CXCR4^-/-^ and plerixafor-treated mice also had less CXCR4-expressing circulating cells and a lower density of peri-bronchial fibrocytes. We demonstrate that targeting CXCR4 has beneficial effects in an animal model of early COPD and provide a framework to translate these preclinical findings to clinical settings in a drug repurposing approach.

**Clinical relevance:** We demonstrate that CXCL12/CXCR4 axis plays an important role in the pathogenesis of early COPD. Inhibition of this axis improves lung function and cardiac tissue remodeling, supporting the future use of CXCR4 inhibitors to slow down the progression of the disease.

## INTRODUCTION

Chronic obstructive pulmonary disease (COPD) is a common disease, characterized by persistent respiratory symptoms and airflow limitation (1). Major risk factors for COPD include chronic exposure to noxious particles, mainly cigarette smoke (CS), and airway infections (2). Early COPD has been recently defined as patients aged <50 years with a smoking exposure >10 pack-years, with a ratio between forced expiratory volume in 1 second (FEV1) and forced vital capacity (FVC) less than 70% or less than the lower limit of normal (LLN), compatible computed tomographic (CT) abnormalities (*i.e*., visual emphysema, air trapping, or bronchial thickening graded mild or worse), and/or evidence of accelerated FEV1 decline of >60 ml/years (3). Twenty-four percent of these early COPD patients developed clinical COPD 10 years later (4). Therapeutic interventions in early COPD patients could help to tackle the disease before reaching irreversible tissue damage (5). However, it is extremely difficult to enrol those patients into trials and collect their bronchial samples, since they are rarely diagnosed, demonstrating the urgent need to develop animal models of early COPD.

Exposure of lung structural and immune cells to CS and infectious agents results in the release of various inflammatory substances, including chemokines (6). The chemokine receptor CXCR4 and its associated ligand CXCL12 appear to be attractive therapeutic targets for early COPD since (i) they are implicated in the migration of inflammatory cells from the blood (7) to the COPD lungs (8), (ii) their expression and function are controlled by other cytokines (9), oxygen concentration (10) and microbial agents (11, 12), which are disrupted in COPD, and (iii) pharmacological blockade of CXCR4 reduces emphysema development in a long-term COPD murine model (13). CXCR4 is notably expressed by fibrocytes, a rare population of circulating fibroblast-like cells (14), that are assumed to play a crucial role in COPD (7, 8).

In the present study, we thus hypothesized that the CXCL12/CXCR4 axis plays a role in the pathological processes leading to COPD development during early adulthood and, hence, that inhibiting CXCR4 would protect from CS-induced airflow limitation and tissue remodeling. In early COPD patients’ samples, as well as in murine model of early COPD, both CXCR4-expressing cells in the blood and CXCL12 in the lung were overexpressed. Using either transgenic mice deficient for CXCR4 or the CXCR4 antagonist plerixafor, we showed that lungs were protected against CS-induced both lung function alteration and fibrocytes accumulation, and that the right heart was resistant to cardiac remodeling.

Some of these findings have been previously reported in abstract form (15, 16).

## METHODS

Complete details concerning methods appear in an online data supplement.

### Study Population

Human lung tissues were obtained from both the “Fibrochir” study (NCT01692444) (8) and “TUBE” (i.e., TissUs Bronchiques et PulmonairEs, sponsored by the University hospital of Bordeaux) biological collection (Table S1). Blood samples of COPD patients were obtained from “COBRA” (i.e., COhort of BRonchial obstruction and Asthma; sponsored by INSERM) (Table S2) and. All subjects gave their written informed consent to participate to the studies. The studies received approval from the local or national ethics committees.

### Dataset transcriptomic analysis

Transcriptomes obtained on human blood and lung tissues were downloaded from the NCBI Gene Expression Omnibus (GEO) database (http://www.ncbi.nlm.nih.gov/geo/) using datasets under the accession codes of GSE100153 and GSE76925, respectively.

### Mouse model of early COPD

Male C57BL/6J mice were obtained from Janvier (St Berthevin, France). CXCR4^lox/lox^ and Cre2ERT2 mice were obtained from The Jackson Laboratory (17) and the “Institut Clinique de la Souris” (18), respectively. All animal studies were performed according to European directives for the protection of vertebrate animals. Agreement was obtained from French authorities (numbers A33-063-907 and A33-318-3) and all the protocols were approved by the local ethics committee.

Ten-week-old mice were exposed for 10 weeks to either room air (RA) or CS from non-filtered research cigarettes (2R4; University of Kentucky, Lexington, KY) using the smoking apparatus (Anitech, Paris, France), as described previously by Almolki *et al*. (19). After 5 weeks of CS exposure, mice were anesthetized and 50 μg of poly(I:C) diluted in PBS, mimicking responses by RNA virus infections (20), or its vehicle control, were administered twice per week for 5 weeks via nasal aspiration. Electrocardiograms (ECG) and Magnetic Resonance Imaging (MRI) were performed during the 10^th^ week of the protocol. After lung function measurement, right ventricular systolic pressure (RVSP) was determined invasively. Following bronchoalveolar lavage (BAL), mice were sacrificed and lung, heart and blood were collected. The Fulton index was calculated as the ratio between the right ventricle (RV) and the left ventricle plus septum (LV + S).

CXCR4 genetic deletion was induced by tamoxifen. CXCR4 pharmacological inhibition was induced by subcutaneous injection of 1 mg/kg plerixafor (Sigma-Aldrich, Saint-Quentin-Fallavier, France), 5 times/week during the last 5 weeks of the exposition protocol.

### Experimental procedures in both human and mice

CXCR4-expressing cells were quantified in the blood and in the lungs by flow cytometry and immunohistochemistry, as described previously (8). CXCL12 and CXCR4 levels were evaluated by ELISA, western blotting and RT-PCR. Proteomics analyses were performed using mass spectrometry, as described previously (21).

### Statistical Analysis

Statistical significance, defined as P < 0.05, was assessed by t-tests and MANOVA for variables with parametric distribution, and by Kruskal-Wallis with multiple comparison z tests, Mann-Whitney tests, Wilcoxon tests and Spearman correlation coefficients for variables with non-parametric distribution.

## RESULTS

### CXCL12 and CXCR4 expression is increased in lung and blood, respectively, in patients with COPD

First, we determined whether the expression of CXCL12 and CXCR4 was altered in patients with COPD by interrogating RNAseq data from lung of control subjects and COPD patients (GSE76925 data set, http://www.ncbi.nlm.nih.gov/geo/). We found that CXCL12 expression was significantly increased in the lung of patients with COPD (GSE76925 data set, figure 1A). In a separate cohort of patients with and without early COPD (table S1), CXCL12 was predominantly localized to bronchial epithelial cells and peri-bronchial infiltrating cells, which are likely immune cells (figure 1B). The CXCL12 surface immunostaining was significantly increased in lungs of patients with early COPD in comparison to control lungs (figure 1B-C). CXCR4 level was not significantly altered in the lungs of patients with COPD compared to those of control smokers at the mRNA level (GSE76925 data set, figure 1D), which was corroborated by a similar result at the protein level by quantification of immunostaining (figure 1E) in patients with and without early COPD. In the blood, CXCR4 was significantly increased, at the mRNA level, in patients with COPD compared to control subjects (figure S1A). In a separate cohort of moderate to mild COPD patients (GOLD I to II, as defined by (1), table S2), the percentage of CXCR4-expressing cells was also increased in the blood of moderate COPD patients in comparison with mild COPD patients (figure S1B-C).

**Figure 1:**
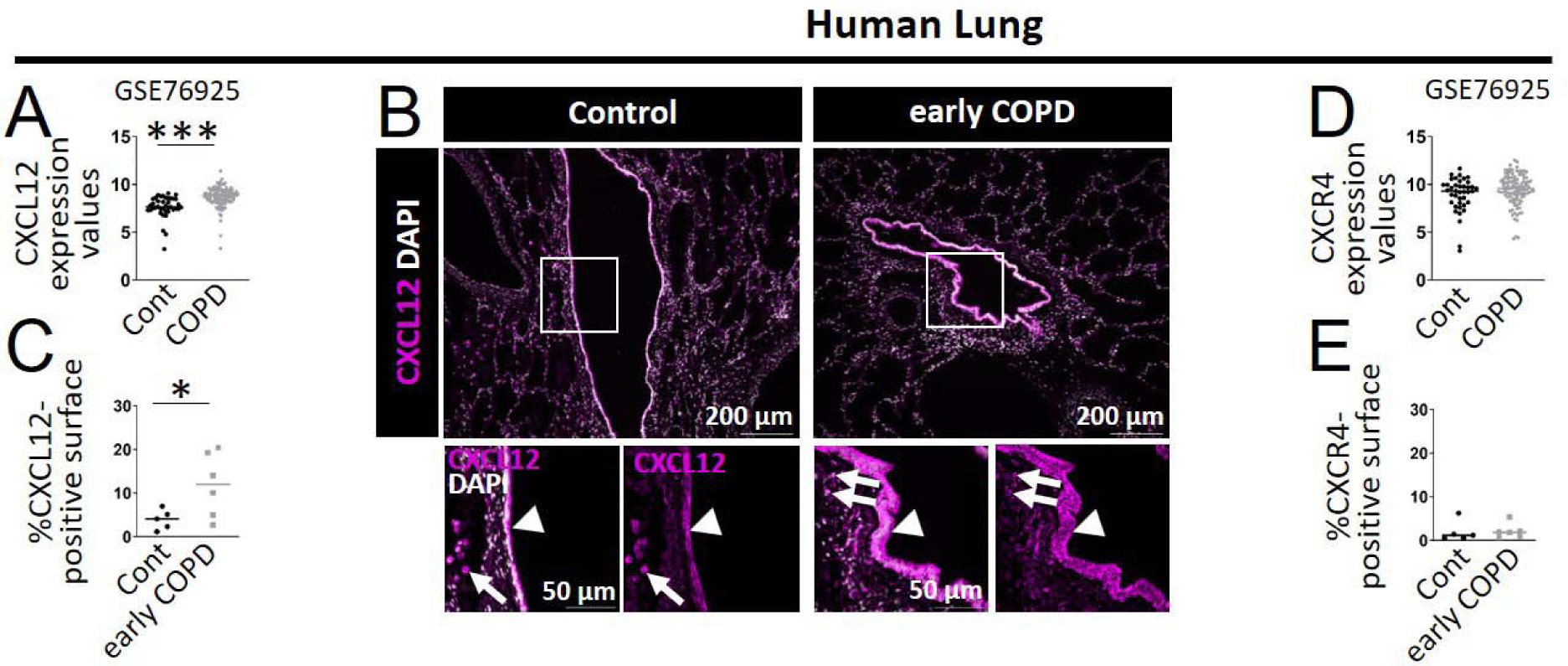
Characterization of CXCR4 and CXCL12 expression in lung of patients with or without COPD. (A, D) CXCL12 (A) and CXCR4 (D) mRNA expression in human lungs. Data are derived from a publicly available GSE-set (GSE76925). Control subjects n=40; patients with COPD n=111. (B) Representative staining of CXCL12 (magenta) and nucleus (white) in peri-bronchial areas of human lungs. Lower panels: higher magnification of the images surrounded by white squares in the upper panels. Images obtained in control *vs* early COPD lungs were acquired with the same exposure time and gain. The white arrowheads and arrows indicate respectively CXCL12-expressing bronchial cells and immune-type cells. (C, E) Quantification of the percentage of positive CXCL12 (C) or CXCR4 (E) surface (normalized by the total area) (n=5 control subjects, n=6 patients with early COPD). Medians are represented as horizontal lines. (A, C, D, E) *: P <0.05, ***: P<0.001, t-test or Mann-Whitney test.

### Chronic CS and poly(I:C) exposure induces functional obstruction, lung inflammation and both lung and heart tissue remodeling in mice

To model early COPD, we used mice at the beginning of adulthood (*i.e*., 10-week old) (22), which were exposed to a combination of CS exposure and intranasal instillations of poly(I:C) in order to mirror the effects of smoking and exacerbations (23). The protocol was based on a previous study (20), where mice were exposed during 5 weeks to cigarette smoke, with poly-IC instillations during the 2 last weeks, based on the intricate relationships between innate immune response and COPD pathogenesis. We reproduced a similar protocol and found no modification of lung function at 5 weeks (figure S2A-D), prompting us to extend to 10 weeks the duration of the protocol. Ten weeks of CS exposure with 10 repeated instillations of poly(I:C) during the last 5 weeks significantly decreased the FEV0.05/FVC ratio (figure 2A-C). Other lung function parameters, in particular respiratory system compliance and tissue elastance, were not modified by CS and poly(I:C) exposure (figure S2E-J). At the end of the protocol, the weight gain of the exposed mice was significantly different from that of the control mice (figure S3A-B). CS and poly(I:C) exposure increased total cell count in the BAL fluid, as well as the absolute neutrophil, lymphocyte and macrophage numbers, 1 day after the last poly(I:C) instillation (figure 2D-F). Although lower, the increase in neutrophils and lymphocytes persisted 4 days after the last poly(I:C) instillation (figure 2D-F). CS and poly(I:C) synergistically enhanced airflow obstruction and BAL inflammation (figure S4). We also analyzed lung proteomics obtained 1 day after the last poly-IC instillation. Of the 137 pathways that were significantly different in early COPD vs control mice, largest upregulated genes were those of the hypercytokinemia/hyperchemokinemia box and of the pathogenesis of influenza and interferon (IFN) signaling (figure 2G-H). In contrast, inhibited pathways were the wound healing and the xenobiotic metabolism PXR signaling pathways (figure 2G-H). Exposed mice had a moderate but significant increase in peri-bronchial fibrosis compared to control mice (figure 2I-J), mimicking the thickening of distal airway tissue in COPD patients (24).

**Figure 2:**
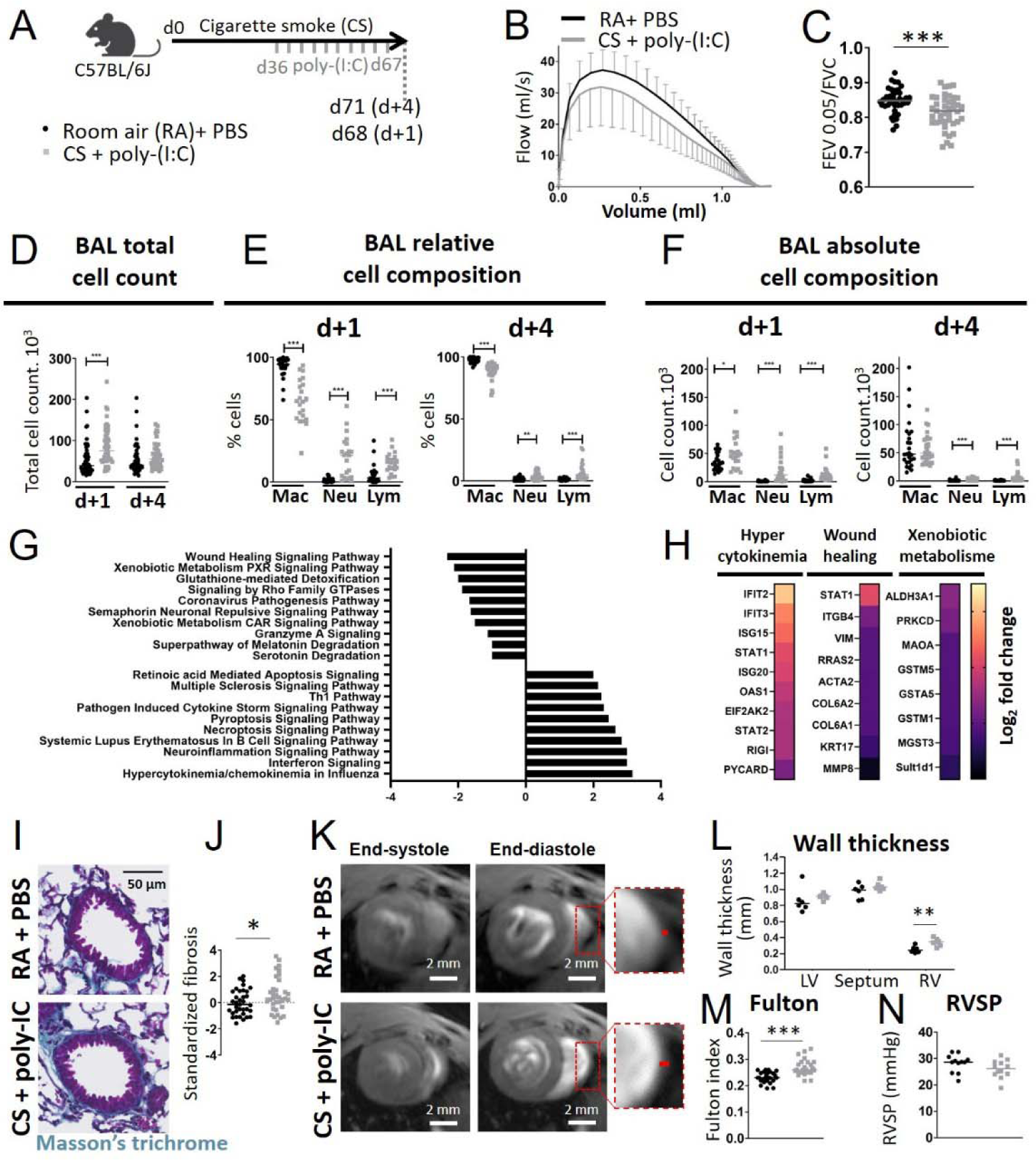
Evaluation of lung function, inflammation, bronchial and heart remodeling in experimental COPD. (A) Mice are exposed either to room air (RA) and challenged with PBS, or exposed to cigarette smoke (CS) and challenged with poly(I:C), during 10 weeks. Mice are sacrificed at day 68 or 71 (*i.e*., 1 day or 4 days after the last poly(I:C) instillation). (B) Average expiratory flow-volume curves of RA+PBS-exposed mice (black curve) and CS+poly(I:C)-exposed mice (grey curve). Lower and upper error bars represent standard deviations for CS+poly(I:C) and RA+PBS-exposed mice, respectively. (C) FEV0.05/FVC (FEV0.05: Forced Expiratory Volume during 0.05 s, FVC: Forced Vital Capacity) of RA+PBS-exposed mice (black circles, n=40) and CS+poly(I:C)-exposed mice (grey squares, n=41). Data represent individual mice. (D) Total cells count in bronchoalveolar lavage (BAL). Kruskal-Wallis test followed by Dunn’s post-tests. (E-F), BAL differential cell recovery 1 day (d+1, RA+PBS: n=24, CS+poly(I:C): n=22) and 4 days (d+4, RA+PBS: n=26, CS+poly(I:C): n=31) after the last poly(I:C) instillation. (G) Top Canonical Ingenuity Pathways significantly altered in CS+poly(I:C) (n=5) *vs* RA+PBS (n=5)-exposed lungs (n=5) obtained 1 day after the last poly-IC instillation, ranked by Z-score (negative and positive for the pathways represented respectively on the top and bottom part of the graph), obtained by Gene Set Enrichment Analysis. (H) Heatmaps of differentially regulated proteins in CS+poly(I:C) vs RA+PBS-exposed lungs, from the pathways “hypercytokinemia/hyperchemokinemia in the pathogenesis of influenza” (left), “wound healing signaling pathway” (middle) and “xenobiotic metabolism PXR signaling pathway” (right). The colour scale indicates the log_2_ fold changes of abundance for each protein. (I) Representative Masson’s trichrome staining to assess peri-bronchial fibrosis. (J) Standardized fibrosis, defined as (peri-bronchial fibrosis (“PF”, %) - mean PFcontrol)/standard deviation PFcontrol). The peri-bronchial fibrosis (“PF”, percentage) is defined by the ratio between the area of segmented pixels in the area of analysis divided by the total peri-bronchial area. RA+PBS: n=35, CS+poly(I:C): n=34. (K) Representative short-axis cine magnetic resonance images of heart at end-systolic stage (left panels) and end-diastolic stage (middle panels) in control mice (top panels) and CS and poly(I:C)-exposed mice (bottom panels). Right panels: higher magnification of RV wall. (L) Wall thickness. RA+PBS: n=6, CS+poly(I:C): n=6. (M) Fulton index, defined as (right ventricle (RV)/ left ventricle plus septum (LV + S). RA+PBS: n=26, CS+poly(I:C): n=24. (N) Right ventricular systolic pressure (RVSP). RA+PBS: n=12, CS+poly(I:C): n=12. (C, E, F, I, L, M, N) Unpaired t test or Mann-Whitney test. *: P<0.05, **: P<0.01, ***: P<0.001.

Then, we investigated the impact of CS and poly(I:C) exposure on cardiac function. There was no evidence of heart failure in our model (figure S5A-B). Moreover, there was no sign of heart rhythm disorder as assessed by ECG monitoring (figure S5C and figure S6A-F). Using MRI, we also found that CS and poly(I:C)-exposition did not alter RV or LV volumes (figure S5D-F). However, we detected a significant increase of RV wall thickness in exposed mice, indicating RV hypertrophy (figure 2K-L). This finding was confirmed by the measurement of the Fulton index (figure 2M). Of note, the RV systolic pressure (RVSP) did not differ between early COPD and control mice (figure 2N).

### CXCR4 and CXCL12 expression is increased in blood and lung, respectively, in experimental COPD

The percentage of CXCR4-expressing cells was significantly increased in the blood of CS and poly(I:C)-exposed mice compared to control mice, 1 day after the last poly(I:C) instillation (figure 3A-B), indicating that experimental COPD replicated the increased levels of CXCR4 in the blood of COPD patients. This effect appeared to be due to a synergy between CS and poly(I:C) (figure S7). The level of CXCR4^+^ circulating cells decreased and became similar between control and exposed mice 4 days after the last poly(I:C) instillation (figure 3B). No significant modification of CXCL12 plasma concentration was evidenced in mice (figure 3C). The percentage of CXCR4-expressing cells was not significantly altered in whole lung homogenates in experimental early COPD (figure 3D-E). In lung homogenates, CS and poly(I:C)-exposure significantly increased CXCL12 protein levels compared to control mice, 1 and 4 days after the last poly(I:C) instillation (figure 3F). In good agreement with our findings in patients’ lungs, CXCL12 was expressed by bronchial epithelial cells and peri-bronchial infiltrating cells (figure 3G), with a higher expression in lungs of animals exposed to CS and poly-(I:C) in comparison to control lungs (figure 3H).

**Figure 3:**
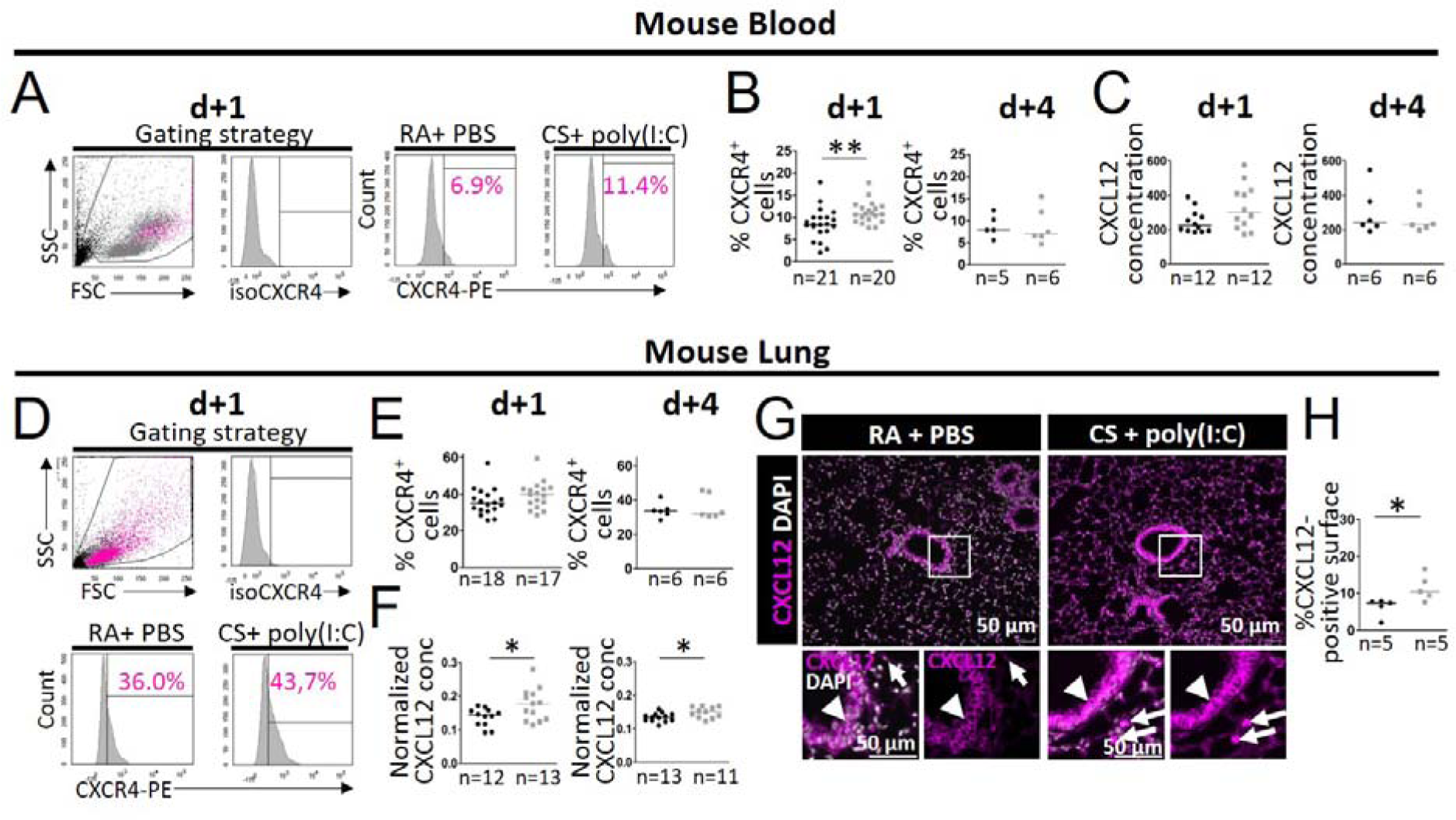
Characterization of CXCR4 and CXCL12 expression in the blood and lung in experimental COPD. Mice are exposed either to room air (RA) and challenged with PBS, or exposed to cigarette smoke (CS) and challenged with poly(I:C), during 10 weeks. They are sacrificed at day 68 or 71 (i.e. 1 day or 4 days after the last poly(I:C) instillation). (A, D) Upper left panels: dot plots represent representative side scatter (SSC, y-axis)-forward scatter (FSC, x-axis) graphs of circulating (A) and lung (D) cells. DAPI^-^ cells, and DAPI^-^ CXCR4^+^ cells are shown respectively in grey and pink. Upper right panels and bottom panels: histograms represent representative cell count (y-axis) versus Phycoerythrin (PE) fluorescence (x-axis) in DAPI^-^ circulating (A) and lung (D) cells. Percentages of CXCR4^+^ cells in the DAPI^-^ cells are shown in pink. (B, E) Levels of circulating (B) and lung (E) CXCR4^+^ cells 1 day (d+1) and 4 days (d+4) after the last poly(I:C) instillation. (C, F) Plasma (C) and lung (F) CXCL12 concentration at d+1 and d+4. For lung CXCL12 concentration, the value is normalized to total protein concentration. (G) Representative staining of CXCL12 (magenta) and nucleus (white) in peri-bronchial areas of mouse lungs. Lower panels: higher magnification of the images surrounded by white squares in the upper panels. Images obtained in RA and PBS *vs* CS and poly(I:C)-exposed have been acquired with the same exposure time and gain. The white arrowheads and arrows indicate respectively CXCL12-expressing bronchial cells and immune-type cells. (H) Quantification of the percentage of positive CXCL12 surface (normalized by the total area). (B, C, E, F, H) Medians are represented as horizontal lines. *: P <0.05, **: P<0.01, Mann-Whitney test.

### Fibrocytes and CXCR4^+^ fibrocytes are increased in lungs of experimental early COPD

To quantify fibrocytes in control and experimental COPD lungs, single cell suspension was stained with CD45 and FSP1 as well as CXCR4. We defined fibrocytes as cells expressing CD45 and a high level of FSP1 (“CD45^+^ FSP1^high^”) (25, 26), which is also in agreement with previous studies performed in human lung (8, 27). The percentage of fibrocytes was significantly increased in CS and poly(I:C)-exposed lungs 1 day after the last poly(I:C) instillation (figure 4A-C). The average percentage of leukocytes (CD45^+^ cells) in control and exposed lungs was 69.2 % and 74.0 %, respectively (p=0.07, figure 4D). In control and experimental early COPD, a vast majority of lung fibrocytes expressed CXCR4 (96.1 and 96.6 %, respectively, figure 4B) and the percentage of CXCR4^+^ fibrocytes was significantly increased in the lungs of exposed mice (figure 4C). Four days after the last poly(I:C) instillation, the total level of fibrocytes decreased and became similar between control and experimental early COPD lungs (figure 4C). Since the flow cytometry approach limits our analysis to the whole lung, we co-stained FSP1 with CD45 in murine lungs, 4 days after the last poly(I:C) instillation (figure 4F) using immunohistochemistry. Similarly to that previously reported in human lung (8), the density of peri-bronchial fibrocytes was higher in CS and poly(I:C)-exposed murine lungs than in control lungs (figure 4G). Moreover, we also observed *in situ* CXCR4-expressing fibrocytes in peri-bronchial areas of human lung samples (“Fibrochir” study (8), figure 4H). Of note, fibrocytes were found in right and left ventricles, but their densities were unchanged by CS and poly(I:C) exposure (figure S8).

**Figure 4:**
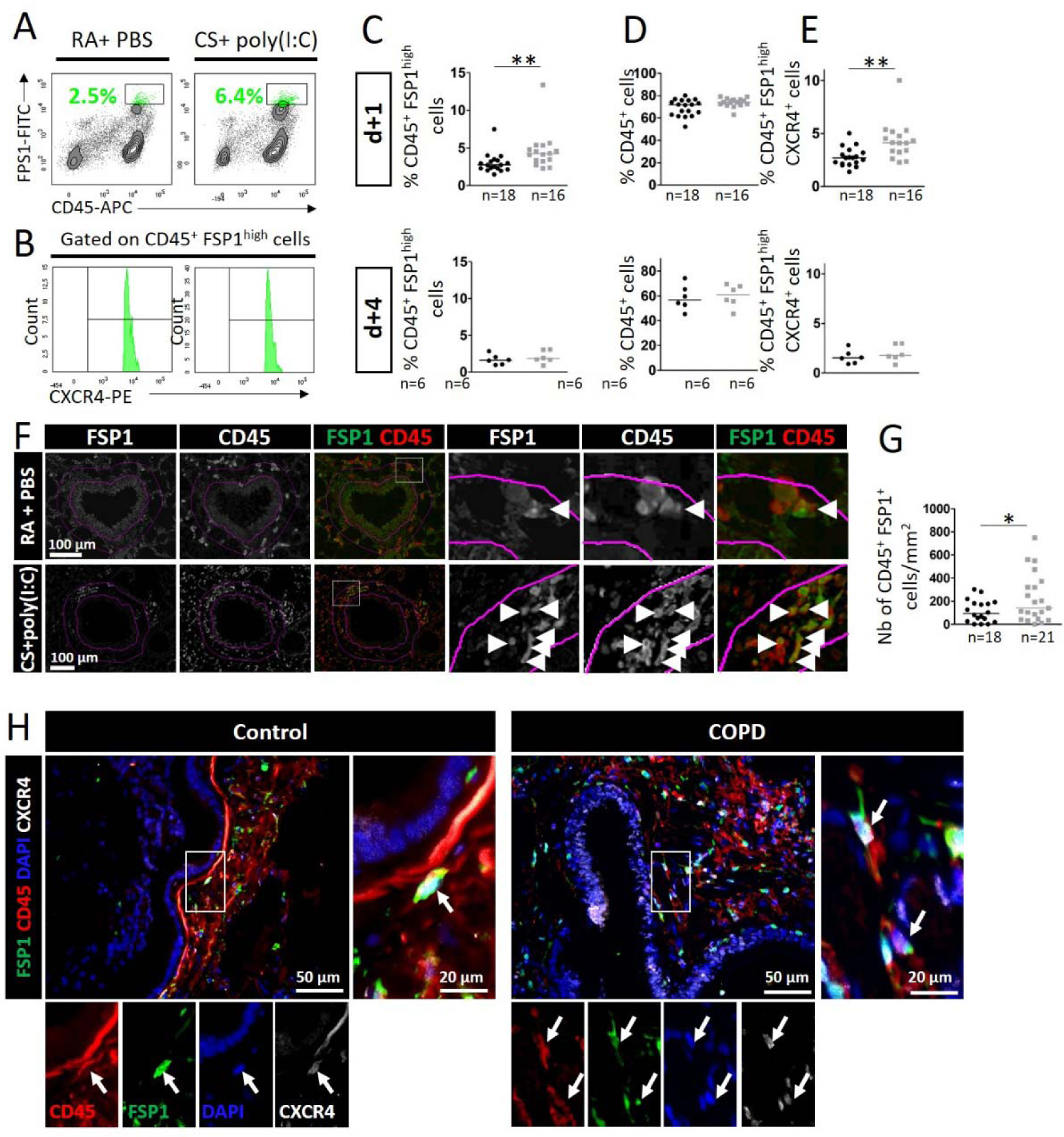
Characterization of lung fibrocyte level and localization in experimental COPD and in patients with or without COPD. (A-G) Mice are exposed either to room air (RA) and challenged with PBS, or exposed to cigarette smoke (CS) and challenged with poly(I:C), during 10 weeks. Mice are sacrificed 1 day (“d+1”) or 4 days (“d+4”) after the last poly(I:C) instillation. a) Representative flow cytometry contour plots in each condition. The x-axis is for CD45-Allophycocyanin (APC) fluorescence and the y-axis, FSP1-Fluorescein-5-isothiocyanate (FITC) fluorescence. Percentages of CD45^+^ FSP1^high^ cells are shown in green. (B) Representative cell count (y-axis) versus CXCR4-Phycoerythrin (PE) fluorescence (x-axis) in CD45^+^ FSP1^high^-cell subsets in mouse lungs. (C-E) Levels of lung CD45^+^ FSP1^high^ cells (C), CD45^+^ cells (D), CD45^+^ FSP1^high^ CXCR4^+^ cells (E), in RA+PBS-exposed mice (black circles) and CS+poly(I:C)-exposed mice (grey squares), at d+1 and d+4. (F) Representative staining of CD45 (green) and FSP1 (red) in peri-bronchial areas (delimited in pink) of lung mice at d+4. The white arrows indicate fibrocytes, defined as CD45^+^ FSP1^+^ cells. (G) Quantification of fibrocyte density (normalized by peri-bronchial area). (H) Representative staining of CD45 (green) and FSP1 (red) in peri-bronchial area. The white arrowheads indicate fibrocytes, defined as CD45^+^ FSP1^+^ cells. (F) Representative staining of CD45 (green), FSP1 (red), nuclei (DAPI, blue) and CXCR4 (white) in peri-bronchial areas of lung samples of control subjects and patients with COPD. The white arrows indicate CXCR4-expressing fibrocytes, defined as CD45^+^ FSP1^+^ CXCR4^+^ cells. (F, H) Left and lower panels: higher magnification of the images surrounded by white squares in the right upper panels. (C-E, G) Medians are represented as horizontal lines. *: P <0.05, **: P<0.01, Mann-Whitney test.

### CXCR4 deletion protects against functional airway obstruction, lung and heart tissue remodeling, and inhibits fibrocyte recruitment into the lungs in experimental early COPD

As mice genetically deficient for CXCR4 die perinatally (28), we generated conditional CXCR4 mutants using floxed allele (17) and a Cre2ERT2 strain (18), thereby obtaining mutants harboring a specific inactivation of CXCR4 at early adulthood after tamoxifen treatment. Intraperitoneal tamoxifen administration, for 5 consecutive days, in CXCR4^lox/lox^/Cre2ERT2 mice was not sufficient to induce a large CXCR4 downregulation, particularly, in the lungs (figure S9A-B). Therefore, we combined intraperitoneal tamoxifen administration and feeding mice with tamoxifen-containing food to obtain a more robust decrease in CXCR4 expression in the lung and bone marrow (figure S9C-E). As tamoxifen-containing food reduced weight gain in mice (figure S3C-D), we used CXCR4^lox/lox^ mice as control mice, and gave all transgenic mice tamoxifen-containing food.

CXCR4^lox/lox^/Cre2ERT2 and CXCR4^lox/lox^ mice were exposed to CS and poly(I:C) for 10 weeks (figure 5A). As expected, CXCR4 expression decreases in the lung and blood of CXCR4^lox/lox^/Cre2ERT2 mice (figure S10). Moreover, FEV0.05/FVC ratio was higher in CXCR4^-/-^ mice compared to CXCR4^lox/lox^ mice, when exposed to CS and poly(I:C) (figure 5B). In the BAL fluid, CXCR4 deletion induced a significant decrease of macrophage and increase of neutrophil percentages, 1 day after the last poly(I:C) instillation (figure 5C-E). CXCR4^-/-^ mice had a significant reduction in collagen deposition around small airways, compared to CS and poly(I:C)-exposed CXCR4^+/+^ mice (figure 5F-G), confirming an important role of CXCR4 in CS-induced bronchial remodeling.

**Figure 5:**
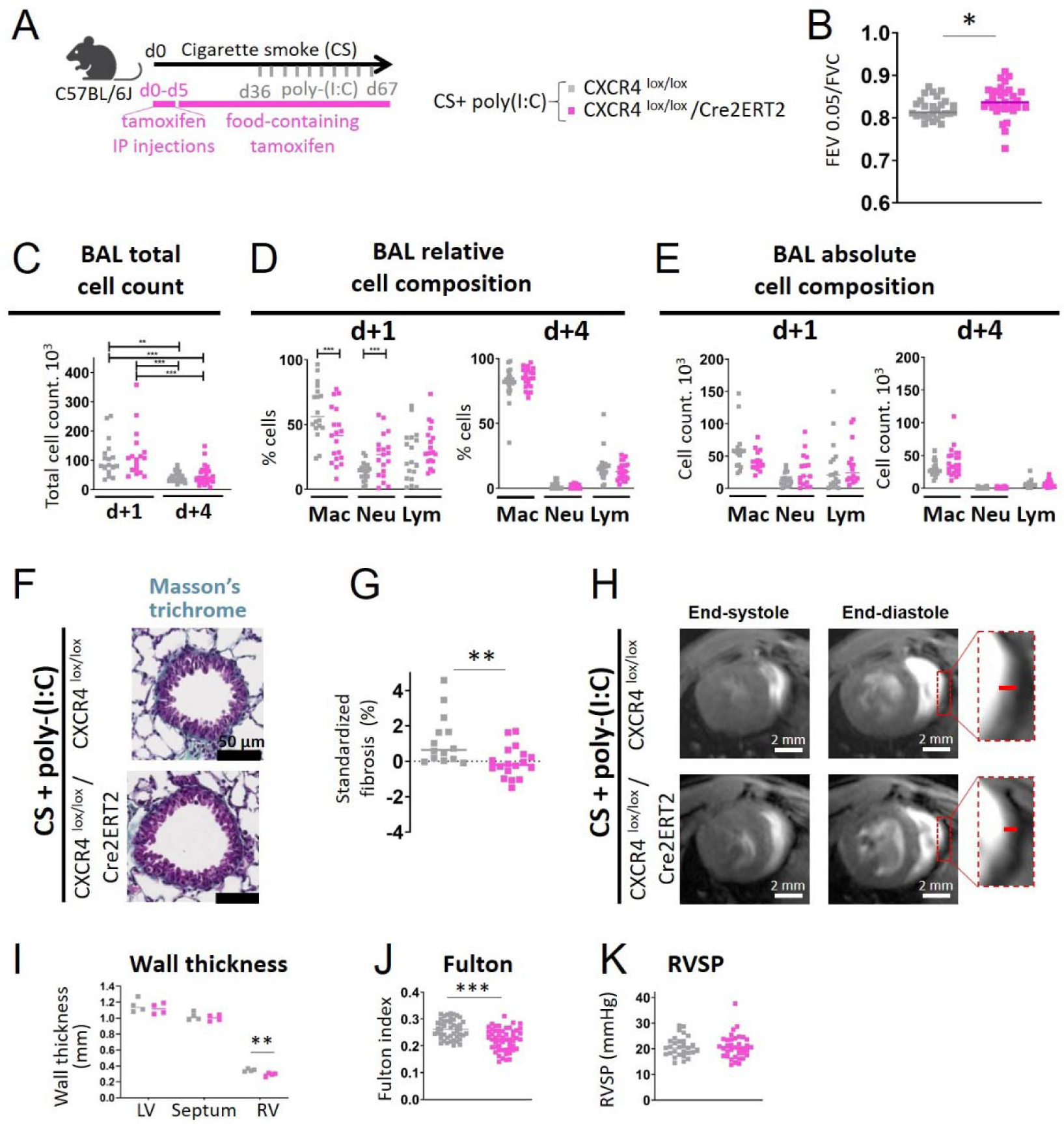
Evaluation of lung function, inflammation, and bronchial and heart remodeling upon conditional knockout of CXCR4 in experimental COPD. (A) CXCR4^lox/lox^ (grey squares) and CXCR4^lox/lox^/Cre2ERT2 (pink squares) mice are exposed to cigarette smoke (CS) and challenged with poly(I:C). All the mice were administered with 1 mg intraperitoneal tamoxifen for 5 consecutive days at the beginning of the COPD protocol, and then fed by tamoxifen-containing food during the remaining protocol. Mice are sacrificed at day 68 or 71 (1 day and 4 days after the last poly(I:C) instillation). (B) FEV0.05/FVC in the different conditions. Data represent individual mice. CXCR4^lox/lox^: n=27, CXCR4^lox/lox^/Cre2ERT2: n=32. (C) Total cells count in bronchoalveolar lavage (BAL). Kruskal-Wallis test followed by Dunn’s post-tests. (D-E) BAL relative (D) and absolute (E) cell composition respectively 1 day and 4 days after the last poly(I:C) instillation. d+1: CXCR4^lox/lox^: n=18, CXCR4^lox/lox^/Cre2ERT2: n=19; d+4: CXCR4^lox/lox^: n=21, CXCR4^lox/lox^/Cre2ERT2: n=23. (F) Representative Masson’s trichrome staining to assess peri-bronchial fibrosis. (G) Standardized fibrosis, defined as (peri-bronchial fibrosis (“PF”, %) - mean PFcontrol)/standard deviation PFcontrol). The peri-bronchial fibrosis (“PF”, percentage) is defined by the ratio between the area of segmented pixels in the area of analysis divided by the total peri-bronchial area. CXCR4^lox/lox^: n=14, CXCR4^lox/lox^/Cre2ERT2: n=18. (H) Representative short-axis cine magnetic resonance images at end-systolic stage (left panels) and end-diastolic stage (middle panels) in CS and poly(IC)-exposed CXCR4^lox/lox^ mice (top panels) and CS and poly(IC)-exposed CXCR4^lox/lox^/Cre2ERT2 mice (bottom panels). Right panels: higher magnification of RV wall. i) Wall thickness. CXCR4^lox/lox^: n=4, CXCR4^lox/lox^/Cre2ERT2: n=4. (J) Fulton index, defined as (right ventricle (RV)/ left ventricle plus septum (LV + S). CXCR4^lox/lox^: n=46, CXCR4^lox/lox^/Cre2ERT2: n=53. (K) Right ventricular systolic pressure (RVSP). CXCR4^lox/lox^: n=29, CXCR4^lox/lox^/Cre2ERT2: n=36. (B, D, E, G, J, K) *: P<0.05, **: P<0.01, ***: P<0.001. Unpaired t test or Mann-Whitney test.

We then assessed the impact of CXCR4 deletion on right heart remodeling. CXCR4 deletion did not affect electrophysiological and hemodynamic (figure S6G-L and figure S11) but it significantly reduced the RV wall thickness as assessed by heart MRI *in vivo* and the Fulton index *ex vivo* in exposed mice (figure 5H-J). RVSP value was unchanged by CXCR4 deletion (figure 5K).

Moreover, we observed reduced flow cytometry levels of CXCR4^+^ fibrocytes in CS and poly(I:C)-exposed whole lungs of CXCR4^-/-^ mice compared with CXCR4^+/+^ exposed mice, 1 day after the last poly(I:C) instillation (figure 6A-D). Four days after the last poly(I:C) instillation, the density of fibrocytes was also significantly decreased around the small airways (figure 6F-G).

**Figure 6:**
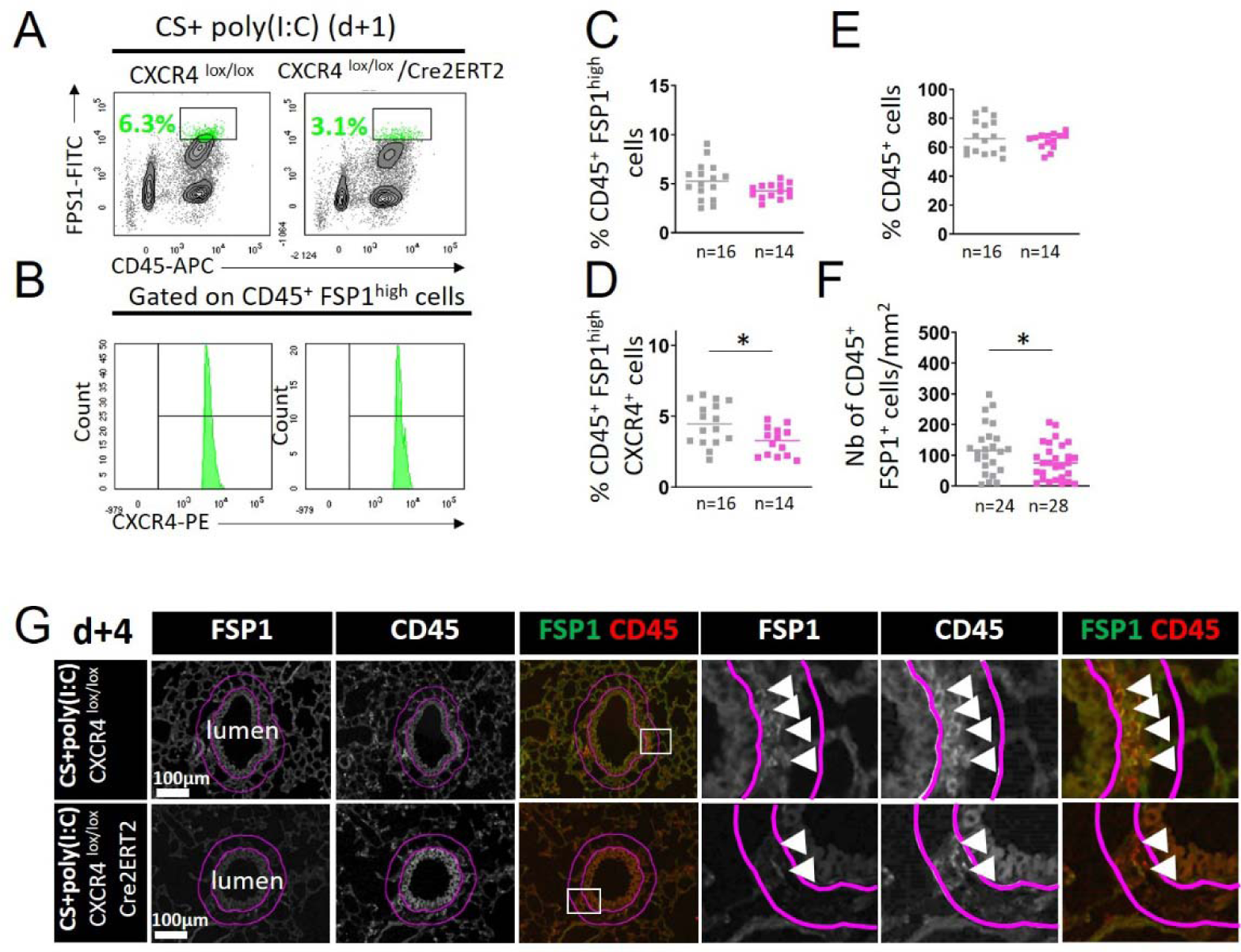
Characterization of fibrocyte level and peri-bronchial density in lung upon conditional knockout of CXCR4 in experimental COPD. CXCR4^lox/lox^ (grey squares) and CXCR4^lox/lox^/Cre2ERT2 (pink squares) mice are exposed to cigarette smoke (CS) and challenged with poly(I:C). Mice are sacrificed at day 68 or 71 (1 day and 4 days after the last poly(I:C) instillation). (A) Representative flow cytometry contour plots in each condition. The x-axis is for CD45-Allophycocyanin (APC) fluorescence and the y-axis, FSP1-Fluorescein-5-isothiocyanate (FITC) fluorescence. Percentages of CD45^+^ FSP1^high^ cells are shown in green. (B) Representative cell count (y-axis) versus CXCR4-Phycoerythrin (PE) fluorescence (x-axis) in CD45^+^ FSP1^high^-cell subsets in mouse lungs. (C-E) Levels of lung CD45^+^ FSP1^high^ cells (C), CD45^+^ FSP1^high^ CXCR4^+^ cells (D), CD45^+^ cells (E) 1 day after the last poly(I:C) instillation in each condition. (F) Quantification of fibrocyte density (normalized by peri-bronchial area) 4 days after the last poly(I:C) instillation in each condition. (G) Representative staining of CD45 (red) and FSP1 (green) in peri-bronchial area (delimited in pink) in each condition. The white arrows indicate fibrocytes, defined as CD45^+^ FSP1^+^ cells. (C-F) Medians are represented as horizontal lines. *: P <0.05, Unpaired t test or Mann-Whitney test.

### Pharmacological inhibition of CXCR4 protects against functional obstruction and inhibits fibrocyte recruitment into the lungs in experimental COPD

We then assessed the efficacy of a CXCR4 antagonist (*i.e*. AMD3100, plerixafor) in the early COPD murine model (figure 7A). Mice weight was not affected by daily subcutaneous plerixafor treatment during the second half of the protocol (figure S3E-F). Plerixafor significantly reduced CXCR4 expression 1 day after the last poly(I:C) instillation in circulating cells, but not in lung cells (figure S12). In exposed-mice treated with plerixafor, the FEV0.05/FVC ratio was significantly higher than vehicle-treated exposed mice (figure 7B). The total number of cells in the BAL fluid and its composition was unchanged by plerixafor treatment (figure 7C-E). The peri-bronchial fibrosis was significantly decreased in the lungs of plerixafor-treated exposed mice (figure 7F-G). To identify pathways that were differentially regulated by the early COPD protocol and plerixafor treatment, we performed proteome analysis on lungs obtained 1 day after the last poly-IC instillation. Comparison analysis revealed a significant inhibition of pathways of inflammation, cell invasion, movement, adhesion, and synthesis of reactive oxygen species (ROS) in relation to plerixafor treatment (figure 7H). These changes were driven by down-regulation of a core set of proteins such as thioredoxin interacting protein (TXNIP), matrix metalloproteinase-8 (MMP-8) and vimentin (figure S13).

**Figure 7:**
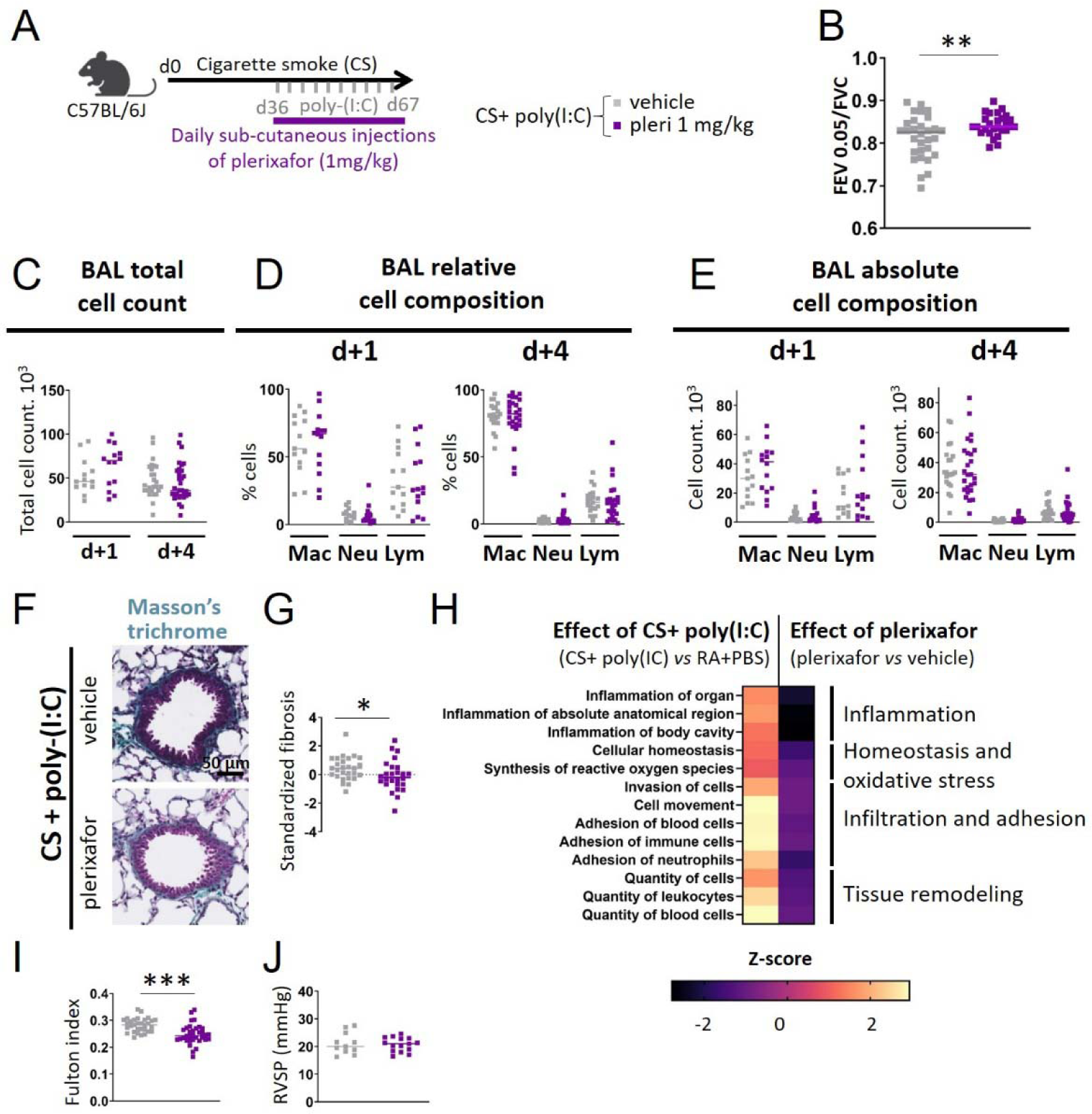
Evaluation of lung function, inflammation, bronchial and heart remodeling upon pharmacological blockage of CXCR4 in experimental COPD. (A) Mice are exposed to cigarette smoke (CS), challenged with poly(I:C) and treated with plerixafor (“pleri”, 1 mg/kg, purple squares) or its vehicle (grey squares). Mice are sacrificed at day 68 or 71 (1 day and 4 days after the last poly(I:C) instillation). (B) FEV0.05/FVC in the different conditions. Data represent individual mice. Vehicle: n=30, pleri: n=27. (C) Total cells count in bronchoalveolar lavage (BAL). Kruskal-Wallis test followed by Dunn’s post-tests. (D-E) BAL relative (D) and absolute (E) cell composition respectively 1 day and 4 days after the last poly(I:C) instillation. d+1: Vehicle: n=13, Pleri: n=14; d+4: Vehicle: n=21, Pleri: n=26. (F) Representative Masson’s trichrome staining to assess peri-bronchial fibrosis. (G) Standardized fibrosis, defined as (peri-bronchial fibrosis (“PF”, %) - mean PFcontrol)/standard deviation PFcontrol). The peri-bronchial fibrosis (“PF”, percentage) is defined by the ratio between the area of segmented pixels in the area of analysis divided by the total peri-bronchial area. Vehicle: n=27, pleri: n=26. (H) Heatmaps of significantly differentially regulated pathways identified by Ingenuity Pathway analysis (IPA; Qiagen), obtained by proteomics comparison of CS+poly(I:C) (n=5) *vs* RA+PBS (n=5)-exposed lungs (n=5) (all treated by vehicle), and plerixafor (n=5) vs vehicle (n=5)-treated lungs (all exposed to CS+poly(I:C)). The colour scale indicates the Z-score value. (I) Fulton index, defined as (right ventricle (RV)/ left ventricle plus septum (LV + S). Vehicle: n=31, pleri: n=34. (J) Right ventricular systolic pressure (RVSP). Vehicle: n=11, pleri: n=15. (B, D, E, G, I, J) *: P<0.05, **: P<0.01, ***: P<0.001. Unpaired t test or Mann-Whitney test.

Treatment with plerixafor also reduced the development of RV hypertrophy in exposed mice (figure 7I). The RVSP, as well as heart electrophysiological properties remained unchanged by CXCR4 inhibition (figure 7J, figure S6M-R).

Flow cytometry 1 day after the last poly(I:C) instillation showed a significant decrease of the numbers of both fibrocytes and CXCR4^+^ fibrocytes in the lungs of exposed mice by plerixafor (figure S14A-E). Four days after the last poly(I:C) instillation, the density of peri-bronchial fibrocytes identified by immunohistochemistry was significantly reduced by plerixafor (figure S14F-G).

## DISCUSSION

Taken together, our data demonstrate that the CXCL12/CXCR4 axis plays a major role in the pathogenesis of lung and heart tissue remodeling of early COPD. Both genetic invalidation and pharmacological inhibition of CXCR4 in our experimental early COPD murine model reduced both the level of CXCR4-positive cells in the peripheral circulation and fibrocyte recruitment into the lungs, along with a proteomic signature consistent with decreased inflammation. As a consequence, CXCR4 inhibition prevented airflow limitation and both lung and cardiac remodeling. Our findings are likely to be translatable to humans as we showed that CXCL12 was increased in the lung of early COPD patients and CXCR4 in the blood of COPD patients.

We designed the present dedicated murine model of early COPD to examine pathophysiological processes at the onset of the disease. Instead of deciphering the relative contribution of CS and poly-(I:C), this study aims at developing and using an early COPD model, based on the following: first, 10-week old mice exposed to CS for 10 weeks would correspond to young human adults chronically exposed to tobacco smoke for several years (22), and poly-(I:C) administration mimics exacerbations occurring in early COPD at a frequency similar to the COPD population (23). Moreover, poly-(I:C) allowed to mimic respiratory viral infections (20), which are more frequent in smokers (29). Second, as in humans, exposed mice exhibited limited but significant airflow limitation together with both neutrophilic/lymphocytic airway inflammation in the BAL, allowing our design to model the bronchial component of COPD (30). Lung histological observations evidenced peribronchial fibrosis, but no obvious signs of alveolar destructions. Neither respiratory system compliance nor tissue elastance was different between control and COPD animals. By contrast, the vast majority of murine COPD models are based on long-term CS exposure (up to 6 months) either mimicking emphysema, or reproducing severe airway obstruction together with emphysema-like changes (31).

The present model also includes features of early RV hypertrophy (32), thus confirming again the relevance of this animal model of early COPD. Cardiovascular diseases are indeed frequently seen in current smokers (33) and patients with COPD (34). Several studies have shown that morphological changes of the right ventricle occur in the initial stages of COPD. Indeed, right ventricular hypertrophy has been described as an early sign of adaptation of the right ventricle to pressure overload in COPD, even in normoxemia or in mild hypoxemia conditions, and initially without affecting right ventricular function (35). Development of right ventricular alterations has also been evidenced very early when airway resistance was increased in preschool-aged children with obstructive respiratory diseases (*i.e*., suffering from chronic coughing and wheezing) (36). Although right ventricular hypertrophy is often described as a consequence of increased pulmonary arterial pressures and development of pulmonary hypertension (PH), right ventricular remodeling has been shown to occur in early stages of COPD before PH development (32, 35, 37). Our mouse model therefore reproduces such early right ventricular alterations regardless of PH development, with significant right ventricular hypertrophy, but without modifications of right ventricular systolic pressure, volume or function. As right ventricular systolic pressure measurements performed in our mice COPD model might not completely reflect PAP, subclinical/minor elevations of the PAP in our model of experimental COPD cannot be ruled out. This hypothesis is in line with the findings of another COPD mice model, where elevations of pressure and higher modification of cardiac remodeling have been found, presumably because of longer time of CS exposure (6 months) (38). More likely, cardiac remodeling could emerge secondary to alternative mechanisms including lung hyperinflation, systemic inflammation, and endothelial dysfunction.

An important result of the present study is the increase of CXCL12 expression in the lungs of early COPD mice and patients, along with an increase in CXCR4^+^-cells in the blood of early COPD mice as well as in COPD patients. The increased level of CXCR4 in both experimental COPD and patients with COPD suggests a therapeutic opportunity to target CXCR4 for COPD treatment. Treating mice with the CXCR4 antagonist plerixafor prevented the increase of CXCR4^+^-cells in the blood, attenuated tissue fibrocytes recruitment and lung function degradation. The beneficial effect of plerixafor on lung function and remodeling in exposed mice, exhibiting increased CXCR4 solely in the blood, was similar with that observed in CXCR4^-/-^ mice. This suggests that such beneficial effects occurred through CXCR4 reduction in the blood rather than in the lungs, which could encourage targeting CXCR4 systemically rather than locally in the lung.

As CXCR4 expression is strongly associated with chemotactic responsiveness (10), CXCR4 up-regulation is likely to result in a dynamic change of cell recruitment into the lungs. In support of this hypothesis, we demonstrated an increased infiltration of fibrocytes into the lungs, that accumulate into peribronchial areas at steady state in CS and poly(I:C)-exposed mice. The elevated lung CXCL12 level in those mice and in patients with early COPD is also likely to participate in fibrocyte chemotaxis from the general circulation to lung tissue, as well as in fibrocyte increase in peribronchial areas, as the binding of CXCL12 on its receptor CXCR4 protects against cell death (39). Pharmacological and genetic targeting of CXCR4 reduced CXCR4-expressing fibrocyte infiltration into the lungs during exacerbations-like events and fibrocyte accumulation around bronchi at the steady state in COPD mice. These results are in agreement with findings obtained in mice treated with another CXCR4 antagonist or with CXCL12 neutralizing antibody, in the context of pulmonary fibrosis, where reduction of fibrocyte accumulation has been found concomitantly of an improvement of lung outcomes (14, 40). Altogether, this indicates that fibrocytes could play a major role in the development of peribronchial fibrosis, either directly through excessive repair processes, or indirectly by paracrine activation of extracellular matrix-producing cells such as fibroblasts and myofibroblasts (41). This could be partially mediated through MMP-8 production, that fibrocyte are able to secrete (42), and that we evidenced as down-regulated by plerixafor treatment along with fibrocyte level reduction. Additionally, and importantly, these fibrocytes could also perpetuate chronic inflammation, through interaction with CD8^+^ T cells. Such cells are indeed implicated in COPD lung pathogenesis (43) and we have recently shown that fibrocytes directly interact with CD8+ T cells in the lungs, leading to increased proliferation and activation (44).

Several other cell types might be implicated in plerixafor-mediated beneficial effects in the lungs, including neutrophils. CXCL12 not only acts as a chemoattractant for neutrophils but also suppresses neutrophils cellular death, allowing other functions for neutrophils such as the production of neutrophil extracellular traps (NETs) (45). CXCR4-expressing neutrophils accumulate in inflamed lungs and they are more prone to produce NETs (46), that may cause tissue damage and contribute to bronchial obstruction. Neutrophils might also control the secondary migration of other cells such as CD8^+^ T cells by leaving CXCL12-containing trails behind during their migration (47). This lead is worthy of investigation and could be the topic of future studies.

Targeting CXCR4 with daily injections of plerixafor during the 5 last weeks of the protocol as well as genetic deletion of CXCR4 decreased smoke induced-right heart remodeling. This extends the findings of another study showing that neutralizing CXCL12 or antagonizing CXCR4 attenuates right ventricular hypertrophy in a rat model of severe pulmonary arterial hypertension (48), and this suggests a general role of the CXCL12-CXCR4 axis in the development of pulmonary hypertension. In the latter study (48), the beneficial effect of CXCL12 neutralization has been attributed to a decrease of vascular pericyte coverage and macrophage infiltration into the lungs, that deserves further investigation in the COPD model. As cardiac comorbidities have a huge impact on the natural history of COPD, with a poor prognosis of COPD patients with pulmonary artery pressure elevations (49), it reinforces the therapeutic interest of CXCR4 as a target for COPD treatment.

Our findings obtained mainly in mice are likely to apply in humans for the following reasons: (i) the expression patterns of CXCL12 are strikingly similar in murine and human lungs, (ii) the CXCL12 and CXCR4 genes originate early in the phylogenetic tree and are highly conserved among vertebrate species, especially between mice and humans (50–52). In the same page, the degree of homology between human and murine CXCR4 and CXCL12 is 90% and 99%, respectively (53). Last, the fact that plerixafor, a drug already used in clinics for CXCR4 antagonization (in the context of hematopoietic stem cells mobilization in the peripheral blood), shows efficacy in our mouse model, further supports the translation of our results to humans.

Collectively, our data show that the CXCL12-CXCR4 axis plays important roles in the pathogenesis of COPD and that inhibitors of the CXCL12-CXCR4 axis could open new therapeutic options in this devastating disease, by improving lung function and decreasing cardiac comorbidities. However, inhibiting the CXCL12-CXCR4 axis may also increase the severity of exacerbations due to its central role in immune response. Therefore, further studies are required to precise the balance between benefits and risks of targeting CXCR4 in COPD.

## ACKNOWLEDGMENTS

We thank the study participants and the staff of the Thoracic Surgery, Radiology, Pathology, Respiratory, Lung Function Testing departments from the University Hospital of Bordeaux (Bordeaux, France), Fabienne Estella for technical assistance with mice protocol, Isabelle Goasdoue, Isabelle Bernis, Natacha Robert, Virginie Niel, and Marine Servat from the clinical investigation center for technical assistance, and Atika Zouine and Vincent Pitard for technical assistance at the Flow cytometry facility (CNRS UMS 3427, INSERM US 005, Univ. Bordeaux, F-33000 Bordeaux, France), Anne-Aurélie Raymond for help with proteomic analysis at the Oncoprot Platform (TBMCore US005). Christel Poujol, Sébastien Marais and Fabrice Cordelières for help with imaging and image analysis et the Bordeaux Imaging Centre (BIC; Bordeaux, France). Microscopy was performed at BIC, a service unit of the CNRS-INSERM and Bordeaux University, a member of the national BioImaging infrastructure of France supported by the French National Research Agency (ANR-10-INBS-04).

The project was supported by a grant from the “Fondation de l’Université de Bordeaux” (Fonds pour les maladies chroniques nécessitant une assistance médico-technique FGLMR/AVAD), a grant from « Aquitaine Science Transfert » (PB) and an unrestricted grant from AstraZeneca, the “Investments in the Future” program managed by the “Agence Nationale de la Recherche” (ANR, grant reference ANR-10-IAHU-04). The COBRA cohort was funded by AstraZeneca, Chiesi, Glaxo-SmithKline, Novartis, Chiesi, Roche and Legs Poix fondation.

## Data sharing

Qualified researchers can request access to all data through the corresponding author.

## Contributors

- conception and design (ID, PB)
- data acquisition (ID, PH, EM, RA, SC, DEH, EE, PE, MC, MD, GC, CB, OO, JWD, TT, VFM)
- data analysis (ID, PH, EM, RA, SC, DEH, RP, MD, GC, CB, JWD, VFM, PB)
- data interpretation (ID, PH, EM, RA, SC, DEH, EE, RP, PE, MD, GC, CB, TT, RM, MZ, VFM, PB)
- resources (PH, MC, MD, MT, HB, POG, MZ, PB)
- drafting the manuscript (ID, PH, PB)
- revision and final approval of the version to be published, agreement to be accountable for all aspects of the work in ensuring that questions related to the accuracy or integrity of any part of the work are appropriately investigated and resolved (All)

## Abbreviations

BAL: Broncho-alveolar lavage
bFGF: basic Fibroblast Growth Factor
BSM: Bronchial smooth muscle
CS: Cigarette smoke
CT: Computed tomographic
ECG: Electrocardiogram
FEV1: Forced expiratory volume in 1 second
FEV0.05: Forced expiratory volume in 0.05 second
FVC: Forced vital capacity
IFN: Interferon
LV: + S Left ventricle plus septum
MRI: Magnetic Resonance Imaging
Poly-(I:C): Polyinosinic–polycytidylic acid
RA: Room air
RV: Right ventricle
RVSP: Right ventricular systolic pressure
VEGF: Vascular Endothelial Growth Factor

