## Supplementary material for "CXCR4 blockade alleviates pulmonary and cardiac outcomes in early COPD": online data supplement

### SUPPLEMENTAL MATERIAL AND METHODS

#### Study Population

Lung tissues for the measurements of CXCL12 and CXCR4 levels were obtained from a separate cohort of patients requiring thoracic lobectomy surgery for nodule or cancer (pN0) (*i.e.*, "TUBE" (*i.e.*, TissUs Bronchiques et PulmonairEs, sponsored by the University hospital of Bordeaux) biological collection, which includes its own local ethic committee (CHUBX 2020/54)). According to the French law and the MR004 regulation, patients received an information form, allowing them to refuse the use of their surgical samples for research. A total of 6 patients with early COPD and 5 control patients (Table S1) were prospectively recruited from the University Hospital of Bordeaux, according to the GOLD guidelines (1).

Human lung tissues for the identification of CXCR4-expressing fibrocytes were obtained from a previously described cohort (2). The study was registered at ClinicalTrials.gov with identifier NCT01692444 ("Fibrochir" study). The study protocol was approved by the research ethics committee ("CPP") and the French National Agency for Medicines and Health Products Safety ("ANSM"). Briefly, subjects more than 40 years of age were eligible for enrolment if they required thoracic lobectomy surgery for cancer (pN0), lung transplantation or lung volume reduction. A total of 17 COPD patients with a clinical diagnosis of COPD according to the GOLD guidelines (1) and 25 non-COPD subjects ("control subjects") with normal lung function testing (*i.e.*, FEV1/FVC > 0.70) and no chronic symptoms (cough or expectoration) were recruited from the University Hospital of Bordeaux.

To quantify the level of CXCR4-expressing cells, blood samples of COPD patients were obtained from "COBRA" (*i.e.*, COhort of BRonchial obstruction and Asthma; sponsored by the French National Institute of Health and Medical Research, INSERM), as outpatients in the Clinical Investigation Centre of the University Hospital of Bordeaux (Table S2).

All subjects gave their written informed consent to participate to the studies. The studies received approval from the local or national ethics committees.

#### Dataset transcriptomic analysis

Transcriptomes obtained on human blood and lung tissues were downloaded from the NCBI Gene Expression Omnibus (GEO) database (http://www.ncbi.nlm.nih.gov/geo/) using datasets under the accession codes of respectively GSE100153 and GSE76925. Differential expression analysis between patients with COPD and control subjects was performed using the GEO2R interactive web tool.

#### Quantification of human circulating CXCR4-expressing cells

Erythrocytes lysis was performed on a 2 ml venous blood sample by adding 5 ml of hypotonic 0.2% NaCl solution during 30s, followed by adding 5 ml of 1.6% NaCl to end with an isotonic solution. This step was repeated if needed, and circulating cells were then washed in cold PBS containing 0.5% bovine serum albumin (BSA, Sigma-Aldrich) and 2 mM Ethylene Diamine Tetra-acetic Acid (EDTA, Invitrogen). Cells were fixed, permeabilized using the IntraPrep Permeabilization Reagent Kit (Beckman Coulter), and stained either with allophycocyanin (APC)-conjugated anti-CXCR4 antibody or with matched IgG2a isotype control (Sony Biotechnology, San Jose, California). Flow cytometry data were acquired using a Canto II 4-Blue 2-Violet 2-Red laser configuration (BD Biosciences). Flow cytometry analysis was performed using Diva 8 (BD Biosciences). Cells were analyzed for CXCR4 expression, with negative control thresholds set using isotype-stained cells. Specific staining for CXCR4 was determined as an increase in positive events over this threshold.

### Animals

Male C57BL/6J mice (~25g, 10 weeks old) were obtained from Janvier (St Berthevin, France). CXCR4$^{lox/lox}$ mice were obtained from the Jackson Laboratory (3). Cre2ERT2 mice were obtained from the Institut Clinique de la Souris (4). CXCR4$^{lox/lox}$/Cre2ERT2 mice were generated by our laboratory by crossing CXCR4$^{lox/lox}$ with Cre2ERT2 mice. The mice were housed in a conventional animal facility with free access to food and water, and environmental enrichment (cylinders). All animal studies were performed according to European and French directives for the protection of vertebrate animals used for Experimental and other Scientific Purposes, and according to the ARRIVE reporting guidelines (5). Agreements were obtained from French authorities (numbers A33-063-907 and A33-318-3) and all the protocols used were approved by the local ethics committee (Comité d’éthique regional d’Aquitaine, protocol numbers: APAFIS#3699-2016012115114794 and APAFIS#20713-2019052014053725).

### Mouse models of early COPD and treatment

Ten-week-old mice were exposed to room air (RA) or cigarette smoke (CS) from non-filtered research cigarettes (2R4; University of Kentucky, Lexington, KY) using the smoking apparatus (Anitech, Paris, France), as described previously by Almolki *et al*. (6). Experiments were performed on at least 15 mice per group. This number was determined using previous studies on lung function (7). No randomisation was used to allocate experimental units to control and treatment groups, but the groups were constituted to have a comparable mean weight between each other. A total of 430 mice was used. The protocol was designed based on those described by Kang *et al*. (8). During the first week, mice received 4 cigarettes twice a day to allow for acclimation (2 cigarettes/session, 2 sessions/day). During the remainder of the exposure, mice received 10 cigarettes per day (5 cigarettes/session, 2 sessions/day). After 5 weeks of CS exposure, mice were anesthetized and 50 µg of poly(I:C) (diluted in 2 volumes of 15 µl PBS) or their control vehicle were administered twice per week (every Monday and Thursday) for 5 weeks via nasal aspiration. Cigarette smoke (CS) exposure was continued during this interval. To minimise confounders, mice from the control group were also introduced in the smoking apparatus, without cigarette smoke exposure, in an alternate manner with CS-exposed mice. There was no expected pain induced by the COPD model. Possible dyspnea could be treated by nebulization of a bronchodilator (salbutamol 1µg/ml). The following endpoints were monitored twice a week: behaviour, external physical appearance (skin and coat condition), breathing, weight. Electrocardiograms (ECGs) and Magnetic Resonance Imaging (MRI) were performed during the $10^{th}$ week of the protocol. At the desired time point, mice were anesthetized with an intraperitoneal injection of 125 mg/kg ketamine and 10 mg/kg xylazine. After lung function measurement, right ventricular systolic pressure (RVSP) was determined by inserting a catheter with fluid-filled force transducers into the right ventricle (RV) on mechanically ventilated mice. Following bronchoalveolar lavage (BAL), mice were sacrificed through injection of 500 mg/kg pentobarbital. Lung, heart, and blood were collected on the same day. The RV and left ventricle plus septum (LV + S) were separated. To assess right ventricular hypertrophy, the Fulton index was calculated as RV / (LV + S) weight.

To induce CXCR4 genetic deletion, dedicated mice were administered with 1 mg intraperitoneal tamoxifen for 5 consecutive days at the beginning of the COPD protocol, and then fed with tamoxifen-containing food during the remaining protocol. For CXCR4 pharmacological inhibition, plerixafor (AMD3100, Sigma-Aldrich) was administrated subcutaneously at a dose of 1 mg/kg body weight, 5 times/week during the last 5 weeks of the exposition protocol.

**ECG recording and analysis in experimental early COPD**

Mice were anesthetized with isoflurane (2-2.5%) and placed on dorsal decubitus position on a heating pad (Minerve, Esternay, France). ECGs were recorded using electrodes in lead II configuration (9). Data acquisition and analysis were carried out using Labchart software (AD-Instruments, Oxford, UK). 1-2 minute(s) of ECG signals recorded at a sampling rate of 200 kHz without variation of heart rate. ECG analysis were conducted as described (10). The heart rate and the P duration as well as the PR, QRS, and QTc intervals, were measured from averaging at least 200 traces. All experiments were realized between 9:00 and 12:00 am.

**MRI acquisition and analysis in experimental early COPD**

MRI acquisitions were performed on a 9.4T Biospec MRI (Bruker, Wissembourg, France) of 20 cm aperture with a transmit- quadrature coil coupled with phase array coil in reception. Animals were anesthetized with 3% isoflurane then maintained at 1.5% (1 $L.min^{-1}$ mixed in air) during acquisitions. The physiological parameters were monitored continuously inside the magnet with ECG electrodes taped on the forepaws and a respiration pad placed on the abdomen (SA Instruments Inc, Stony Brook, USA).

Several scout images were acquired to determine the short axis view of the mouse heart. Short-axis cine IG-FLASH (intragate fast low angle shot) sequence was acquired for each animal with the following parameters: TE/TR= 1.8/51.2 ms; flip angle= 40°; 8 contiguous slices; in-plane resolution= 100x100 $\mu m^2$; slice thickness= 0.8 mm; 20 frames per cardiac cycle; 150 averages; respiratory triggering and retrospective reconstruction with cardiac cycle; acquisition time= 19 min 11 s. These MR images were used to assess: (i) end-diastolic and end-systolic volumes of left ventricle (LV) and right ventricle (RV); (ii) ejection fraction of both ventricles, (iii) cardiac output and (iv) wall thickness of mid-LV, mid-septum and mid-RV at end-diastolic stage. Images were analyzed on Horos (LGPL-3.0) with MRHeart plugin (11).

**Lung function in experimental early COPD**

Mice were anesthetized with an intraperitoneal injection of 125 mg/kg ketamine and 10 mg/kg xylazine (Centravet, Dinan, France). Tracheostomy was performed using an 18G metal cannula. The animal was then connected to the small animal ventilator (flexiVent, Scirecq) and mechanically ventilated at a respiratory rate of 150 breaths/min, with a tidal volume at 10 mL/kg and a PEEP of 3 cm$H_2O$. Forced oscillation measurements were performed using the single-frequency forced oscillation maneuver ("Snapshot-150 perturbation") and the low frequency forced oscillation maneuver ("Quick Prime-3 perturbation"). Respiratory system resistance (Rrs), respiratory system compliance (Crs), respiratory system elastance (Ers), Newtonian resistance ($R_N$), tissue damping (G) and tissue elastance (H) were calculated from the forced oscillation measurements. The NPFE maneuver was then performed by inflating the mouse lungs to a pressure of 30 cm $H_2O$ over 1 second, hold this pressure for 2 seconds before connecting the animal's airways to the negative pressure reservoir (-50 cm $H_2O$) for 2 seconds. The forced expired volume over 0.05 second (FEV0.05), forced vital capacity (FVC) and the peak expiratory flow (PEF) were calculated directly from the flow-volume loop generated during lung deflation. In every mouse, each maneuver was repeated until at least 2 acceptable measurements (coefficient of determination ≥ 0.95) were recorded. The median of acceptable measurements was calculated. If 2 acceptable measurements were not obtained, the measurements were excluded.

**Bronchoalveolar lavage (BAL) in experimental early COPD**

BAL was performed by perfusing through the cannula with two aliquots of 0.3 ml sterile saline. Total cell counting was performed using Neubauer chamber. Cytospins (Fisher Scientific, Illkirch, France) with Diff-Quick (RAL Diagnostics, Martillac, France) staining were made for differential cell counting (at least 180 cells) that is evaluated in a blinded manner. If less than 180 cells were counted or if the BAL was contaminated with blood, the sample was excluded. The cellular contents and BAL fluid were separated by centrifugation, and the BAL fluid was stored in aliquots at –80°C.

**Lung digest cells purification in experimental early COPD**

The right lungs were dissected out and the post-caval lobes were removed for protein assays. The remaining lobes were cut into 1 $mm^3$ fragments and digested with 40 units/ml collagenase for 45 min at 37°C. Lung cells were purified with a 70-µm cell strainer. Erythrocyte lysis was performed by incubating cell pellet with ACK solution (Ozyme, Saint-Cyr-l'École, France) during 1.5 min.

**Circulating cells purification in experimental early COPD**

Venous blood was taken from the abdominal aorta. Plasma was first separated from the whole blood by centrifugation. Hypotonic lysis of red blood cells was performed on the remaining fraction. This step was repeated if needed.

**Circulating and lung cells characterization by flow cytometry in experimental early COPD**

After purification, lung digest and circulating cells were washed in PBS, stained with 4',6-diamidino-2-phénylindole, dichlorhydrate (DAPI, Fisher Scientific, Illkirch, France), fixed and permeabilized overnight with Foxp3 buffer set (Fisher Scientific). Cells were first stained with biotinylated anti-collagen I (Abcam, Cambridge, UK), anti-FSP1 (Agilent, Les Ulis, France) or matched IgG isotype control, and then stained respectively with Alexa488-conjugated streptavidin or FITC–conjugated anti-rabbit antibody (Fisher Scientific). Next, the cell pellet was incubated with anti-CD45- Allophycocyanin (APC), anti-CXCR4- Phycoerythrin (PE), or isotype controls

(BD Biosciences, San Jose, CA). FACS data were acquired using a Canto II 4-Blue 2-Violet 2-Red laser configuration (BD Biosciences). Flow cytometry analysis was performed using Diva 8 (BD Biosciences).

**Lung tissue preparation for protein assays (ELISA and proteomic)**

The post-caval lobes of murine right lungs and parenchyma samples from human subjects were homogenized using bead-beating homogenizer (Precellys®) in ice-cold radio-immunoprecipitation assay (RIPA) lysis buffer supplemented with 10 µL/mL protease inhibitors (Sigma-Aldrich). Homogenates were centrifuged at 13000 g for 10 min, the supernatants were collected and frozen at –80°C until required.

**CXCL12 dosages**

Lung homogenates and plasma were assayed for CXCL12 levels by enzyme-linked immunosorbent assay (ELISA), according to the manufacturer's instructions (Biotechne, Minneapolis, USA). For lung homogenates, CXCL12 levels were normalized to total tissue protein content, the latter being determined by the Lowry method (Bio-Rad, Hercules, CA, USA).

**Western blot**

Human parenchyma samples were assayed for CXCR4 levels by Western blotting. Total tissue protein extracts were loaded onto a 4-20 % SDS-PAGE gel (Bio-Rad), transferred onto nitrocellulose membrane and revealed with a rabbit anti-CXCR4 antibody (Abcam). HRP-coupled anti-rabbit secondary antibody was used for revelation using ChemiDoc imaging instrument (Bio-Rad). Protein expressions were normalized using total loading protein (Stain-free system, Bio-Rad).

**Quantitative Real-Time PCR**

Total RNAs were extracted from lung or bone marrow using RNAeasy kit, according to the manufacturer's instructions (Qiagen, Hilden, Germany). 100 ng RNA were reverse-transcribed using the High Capacity cDNA reverse transcription kit from Applied Biosystems (Waltham, MA, USA). cDNA samples were then analyzed by qPCR using Quantinova SYBR® Green supermix (Qiagen) through the CFX Connect real-time PCR detection system (Bio-Rad). Mouse CXCR4 and GAPDH primers (CXCR4, Forward 5′- GACTGGCATAGTCGGCAATG-3′ and Reverse 5′-AGAAGGGGAGTGTGATGACAAA -3′ GAPDH, primers: Forward 5′-AGGTCGGTGTGAACGGATTTG-3′, Reverse 5′-TGTAGACCATGTAGTTGAGGTCA-3′) were purchased from Sigma-Aldrich. CXCR4 mRNA expression was determined using the comparative $2^{-\Delta\Delta Ct}$ method and normalized to the mRNA expression level of endogenous reference (GAPDH).

**Quantification of airway remodeling**

The left lungs were embedded in paraffin. Sections of 2.5 µm thick were cut, deparaffinized through three changes of xylene and rehydrated through graded alcohols to water, and stained with Masson's Trichrome stain for peribronchial fibrosis. The sections were imaged using a slide scanner Nanozoomer 2.0HT (Hamamatsu Photonics, Massy, France) using objective UPS APO 20X NA 0.75 combined to an additional lens 1.75X, leading to a final magnification of 35X. Virtual slides were acquired with a TDI-3CCD camera. Brightfield images were acquired with the NDP-scan software (Hamamatsu) and processed with ImageJ software. All tissue area was assessed in a blinded fashion for mice' characteristics. A color threshold was applied to the bright-field image to select the area of peribronchial trichrome staining. Bronchial lumen and basement membrane were manually determined. The area of analysis was defined as an area of a 20-µm greater radius than the basement membrane, without the lumen area. The peribronchial fibrosis ("PF", percentage) was defined by the ratio between the area of segmented pixels in the area of analysis divided by the total peribronchial area. The standardized fibrosis was defined by (PF-mean PFcontrol)/standard deviation PFcontrol).

**Immunohistochemistry**

The sections were stained with mouse anti-CXCL12 (Biotechne) was used for CXCL12 localization in both mouse and human lungs, and rabbit anti-CXCR4 (Abcam) for CXCR4 localization in human lungs. Rabbit anti-FSP1 polyclonal antibody (Dako, Agilent, Santa Clara, USA) and rat anti-CD45 monoclonal antibody (BD Biosciences, San Jose, CA) were used for fibrocytes identification in mouse lungs; for human lungs, mouse anti-human CD45 (clone HI30, Sony) was used. Alexa 647-conjugated mouse anti-CXCR4 (Santa Cruz) was used for CXCR4+ fibrocytes identification in human lungs.

For CXCL12 and CXCR4 simple staining, the sections were re-incubated respectively with Alexa568–conjugated anti-mouse (Fisher Scientific) and Alexa 568-conjugated anti-rabbit antibodies and DAPI for nucleus staining. For CD45-FSP1 double staining and CD45-FSP1-CXCR4 triple staining in human lungs, the sections were re-incubated with Alexa488–conjugated anti-Rabbit and Alexa568–conjugated anti-rat (Fisher Scientific) antibodies and DAPI for nucleus staining. Fluorescent acquisitions were performed as previously described (2), using the slide scanner Nanozoomer with fluorescence imaging module, using a mercury lamp (LX2000 200W - Hamamatsu Photonics) and the set of filters adapted for DAPI, Alexa 488, Alexa 568 and Alexa 647. Fluorescence images were acquired with the NDP-scan software (Hamamatsu) and processed with ImageJ software. Tissue area and cell measurements were all performed in a blinded fashion for human and mice' characteristics.

To quantify the density of $CD45^+$ $FSP1^+$ cells or the surface of CXCR4 or CXCL12 positive staining, a binary threshold was applied to fluorescent images corresponding to CD45, FSP1, CXCR4 and CXCL12 staining. The percentage of positive staining was defined as a ratio between the surface of positive staining and the total surface. Tissue fibrocytes were defined as cells double positive for both cytoplasmic FSP1 and plasma membrane CD45 on the merged threshold images. Bronchial lumen and basement membrane were determined on DAPI staining. As there was no systematic smooth muscle layer to properly delimits the lamina propria, we chose to define the peribronchial area as a 100% enlargement of bronchial lumen. The density of $FSP1^+$ $CD45^+$ cells or $CXCR4^+$ cells was defined by the ratio between the number of double positive cells in the lamina propria divided by the lamina propria area.

**Label-free quantitative proteomics**

5 independent biological replicates on total protein extracts from lungs of mice exposed either to room air, challenged with PBS and treated by plerixafor's vehicle (control condition), or exposed to cigarette smoke, challenged with poly(I:C) and treated with plerixafor or its vehicle. 10 µg of proteins were loaded on a 10% acrylamide SDS-PAGE gel and proteins were visualized by Colloidal Blue staining. Migration was stopped when samples had just entered the resolving gel and the unresolved region of the gel was cut into only one segment. Each SDS-PAGE band was cut into 1 mm x 1 mm gel pieces. Gel pieces were de-stained in 25 mM ammonium bicarbonate ($NH_4HCO_3$), 50% Acetonitrile (ACN) and shrunk in ACN for 10 min. After ACN removal, gel pieces were dried at room temperature. Proteins were first reduced in 10 mM dithiothreitol, 100

mM $NH_4HCO_3$ for 60 min at 56°C then alkylated in 100 mM iodoacetamide, 100 mM $NH_4HCO_3$ for 60 min at room temperature and shrunk in ACN for 10 min. After ACN removal, gel pieces were rehydrated with 50 mM $NH_4HCO_3$ for 10 min at room temperature. Before protein digestion, gel pieces were shrunk in ACN for 10 min and dried at room temperature. Proteins were digested by incubating each gel slice with 10 ng/µl of trypsin (V5111, Promega) in 40 mM $NH_4HCO_3$, rehydrated at 4°C for 10 min, and finally incubated overnight at 37°C. The resulting peptides were extracted from the gel by three steps: a first incubation in 40 mM $NH_4HCO_3$ for 15 min at room temperature and two incubations in 47.5% ACN, 5% formic acid for 15 min at room temperature. The three collected extractions were pooled with the initial digestion supernatant, dried in a SpeedVac, and resuspended with 0.1% formic acid for a final concentration of 0.02 µg/µL. NanoLC-MS/MS analysis were performed using an Ultimate 3000 RSLC Nano-UPHLC system (Thermo Scientific, USA) coupled to a nanospray Orbitrap Fusion™ Lumos™ Tribrid™ Mass Spectrometer (Thermo Fisher Scientific, California, USA). Each peptide extracts were loaded on a 300 µm ID x 5 mm PepMap $C_{18}$ precolumn (Thermo Scientific, USA) at a flow rate of 10 µL/min. After a 3 min desalting step, peptides were separated on a 50 cm EasySpray column (75 µm ID, 2 µm $C_{18}$ beads, 100 Å pore size, ES903, Thermo Fisher Scientific) with a 4-40% linear gradient of solvent B (0.1% formic acid in 80% ACN) in 115 min. The separation flow rate was set at 300 nL/min. The mass spectrometer operated in positive ion mode at a 1.9 kV needle voltage. Data were acquired using Xcalibur 4.4 software in a data-dependent mode. MS scans (m/z 375-1500) were recorded at a resolution of R = 120000 (@ m/z 200), a standard AGC target and an injection time in automatic mode, followed by a top speed duty cycle of up to 3 seconds for MS/MS acquisition. Precursor ions (2 to 7 charge states) were isolated in the quadrupole with a mass window of 1.6 Th and fragmented with HCD@28% normalized collision energy. MS/MS data were acquired in the Orbitrap cell with a resolution of R=30000 (@m/z 200), a standard AGC target and a maximum injection time in automatic mode. Selected precursors were excluded for 60 seconds. Protein identification and Label-Free Quantification (LFQ) were assessed in Proteome Discoverer 2.5. MS Amanda 2.0, Sequest HT and Mascot 2.5 algorithms were used for protein identification in batch mode by searching against a Uniprot Mus musculus database (55 338 entries, release January, 2022). Two missed enzyme cleavages were allowed for the trypsin. Mass tolerances in MS and MS/MS were set to 10 ppm and 0.02 Da. Oxidation (M) and acetylation (K) were searched as dynamic modifications and carbamidomethylation (C) as static modification. Peptide validation was performed using Percolator algorithm and only “high confidence” peptides were retained corresponding to 1% false discovery rate at peptide level (12). Minora feature detector node (LFQ) was used along with the feature mapper and precursor ions quantifier. The normalization parameters were selected as follows: 1: Unique peptides, 2: Precursor abundance based on intensity, 3: Normalization mode: total peptide amount, 4: Protein abundance calculation: summed abundances, 5: Protein ratio calculation: pairwise ratio based, 6: Imputation mode: Low abundance resampling and 7: Hypothesis test: t-test (background based). Quantitative data were considered for master proteins, quantified by a minimum of 2 unique peptides, a fold changes above 2 and a statistical p-value adjusted using Benjamini-Hochberg correction for the FDR lower than 0.05. The mass spectrometry proteomics data have been deposited to the ProteomeXchange Consortium via the PRIDE (13) partner repository with the dataset identifier PXD040105. Proteins were clusterized according to their functions by using the Kyoto Encyclopedia of genes and genome analysis in the search tool for retrieval of interaction between genes and proteins (STRING) database. More global analysis of the data was performed through Ingenuity Pathway Analysis (IPA; Qiagen). We also used the ‘Core Analysis’ package to identify relationships, mechanisms, functions, and pathways relevant to a dataset. Comparative analyses were finally performed with IPA using the ‘Comparative Analysis’ module.

**Statistical Analyses**

Statistical significance, defined as $P < 0.05$, was analyzed by t-tests and MANOVA for variables with parametric distribution, and by Kruskal-Wallis with multiple comparison z tests, Mann-Whitney tests, Wilcoxon tests and Spearman correlation coefficients for variables with non-parametric distribution, with GraphPad Prism 6 software.

**SUPLEMENTAL TABLES**

**Table S1: Patient characteristics (for CXCL12 and CXCR4 localization in human lungs)**

| | **Control** | **Early COPD** | **P-value** |
|---|---|---|---|
| n | 5 | 6 | |
| Age (years) | 47.7 ± 4.8 | 43.2 ± 3.3 | 0.11 |
| Sex (Men/Woman) | 2/3 | 3/3 | >0.99 |
| Current smokers (Y/N) | 2/3 | 4/2 | 0.57 |
| Former smokers (Y/N) | 3/2 | 2/4 | 0.57 |
| Pack years (no.) | 30.0 ± 21.6 | 30.0 ± 14.5 | 0.88 |
| CT abnormalities (Y/N) | 0/5 | 3/3* | 0.18 |
| **PFT** | | | |
| $FEV_1$ (% pred.) | 89.3 ± 13.7 | 76.7 ± 22.3 | 0.72 |
| $FEV_1$/FVC ratio (%) | 80.2 ± 0.05 | 70.2 ± 0.14 | 0.26 |
| FEV1/FVC<0.70 (Y/N) | 0/5 | 3/3 | 0.18 |
| **Treatment** | | | |
| Use of oral corticoids (Y/N) | 0/5 | 0/6 | >0.99 |
| Use of inhaled corticoids (Y/N) | 0/5 | 1/5 | >0.99 |
| Use of LABA/LAMA | 0/5 | 2/4 | 0.45 |

Plus–minus values are means ± SD. CT: computed tomography; PFT, pulmonary function test; FEV1, forced expiratory volume in 1 second; FVC, forced vital capacity; LABA, long-acting beta-agonists; LAMA, long-acting muscarinic antagonists. P-values were calculated with the use of

Fisher’s exact test for comparison of proportions and the Mann–Whitney test for comparison of nonparametric variables.

In early COPD patients, 3 of the 6 patients are defined as early COPD based on a FEV1/FVC<0.70, the 3 other ones show CT abnormalities.

* CT abnormalities: 2 patients with bronchiectasis, 1 patient with emphysematous lesions

**Table S2: Patient characteristics (for human circulating $CXCR4^+$ cells measurements)**

| | COPD GOLD I | COPD GOLD II | P-value |
|---|---|---|---|
| n | 5 | 6 | |
| Age (years) | 70.2 ± 9.1 | 69.2 ± 7.5 | 0.78 |
| Sex (Men/Woman) | 3/2 | 5/1 | 0.54 |
| Current /Former smokers | 2/3 | 1/5 | 0.54 |
| Pack years (no.) | 45.4 ± 10.9 | 51.5 ± 29.8 | 0.85 |
| Smoking duration (years) | 44.8 ± 11.6 | 43.3 ± 7.4 | >0.99 |
| **PFT** | | | |
| $FEV_1$ (% pred.) | 86.2 ± 5.8 | 64.5 ± 10.8 | **0.004** |
| $FEV_1$/FVC ratio (%) | 64.6 ± 2.8 | 59.9 ± 4.2 | **0.04** |
| FVC (% pred.) | 105.4 ± 12.9 | 85.2 ± 20.2 | 0.23 |
| **Six-minute walk test distance (m)** | 460 ± 146 | 410 ± 130 | 0.92 |
| **Arterial blood gases** | | | |
| $PaO_2$ (mm Hg) | 77.1 ± 14.7 | 86.7 ± 6.9 | 0.46 |
| $PaCO_2$ (mm Hg) | 34.7 ± 2.1 | 35.0 ± 5.9 | 0.90 |
| **Treatment** | | | |
| Use of oral corticoids (Y/N) | 0/5 | 0/6 | >0.99 |

| | | | |
|---|---|---|---|
| Use of inhaled corticoids (Y/N) | 0/5 | 3/3 | 0.18 |
| Use of LABA/LAMA | 4/1 | 4/2 | >0.99 |

Plus–minus values are means ± SD. PFT, pulmonary function test; $FEV_1$, forced expiratory volume in 1 second; FVC, forced vital capacity; $PaO_2$, partial arterial oxygen pressure, $PaCO_2$, partial arterial carbon dioxide pressure; LABA, long-acting beta-agonists; LAMA, long-acting muscarinic antagonists. P-values were calculated with the use of Fisher's exact test for comparison of proportions and the Mann–Whitney U-test for comparison of nonparametric variables.

### SUPLEMENTAL FIGURES

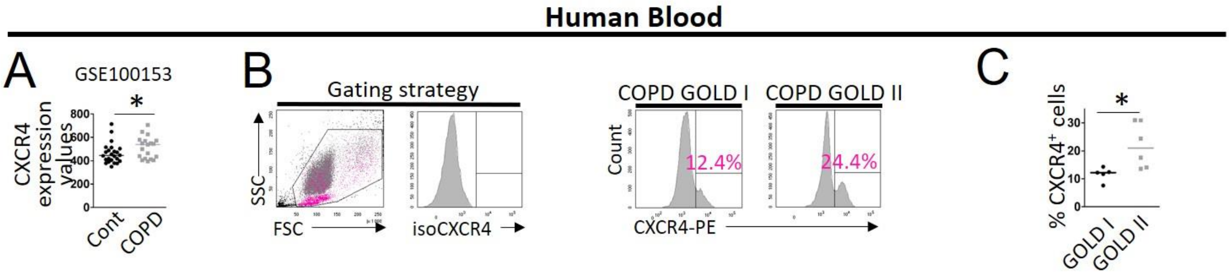


**Figure S1: Characterization of CXCR4 expression in the blood of patients with or without COPD**. (A) CXCR4 mRNA expression in whole blood. Data are derived from a publicly available GSE-set (GSE100153). Control subjects n=24; patients with COPD n=19. (B) Dot plot represents representative side scatter (SSC, y-axis)-forward scatter (FSC, x-axis) graphs of circulating cells. CXCR4$^+$ cells are shown in pink. Histograms represent representative cell count (y-axis) versus Phycoerythrin (PE) fluorescence (x-axis) in circulating cells. Percentages of CXCR4$^+$ cells are indicated in pink. (C) Levels of circulating CXCR4$^+$ cells in patients with moderate COPD (GOLD I, defined by forced expiratory volume in 1 second (FEV1) $\geq$ 80% predicted, n=5) and mild COPD (GOLD II, defined by 50% $\leq$ FEV1 < 80% predicted, n=5). (A, C) Medians are represented as horizontal lines. *: P <0.05, Mann-Whitney test.

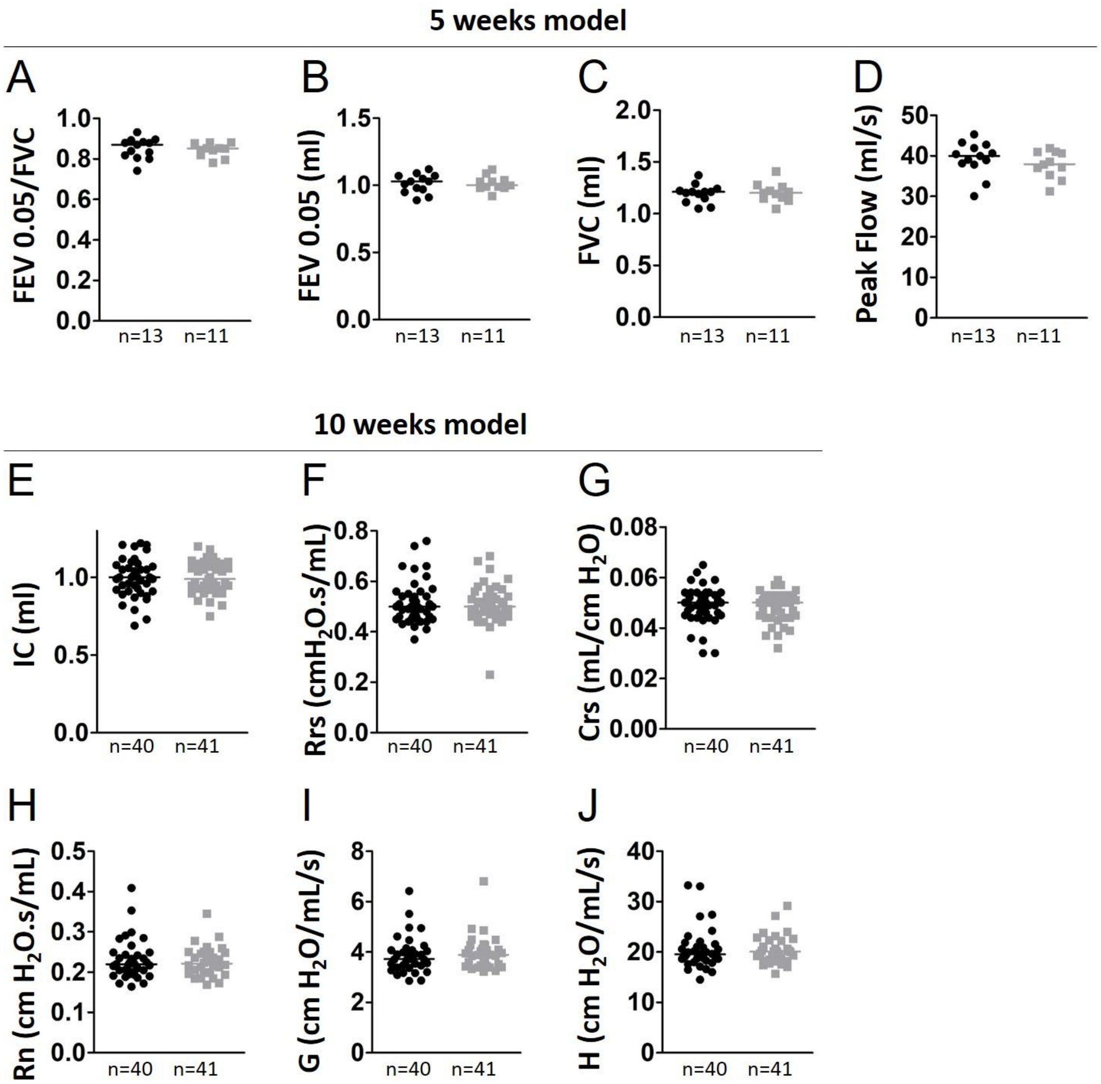


**Figure S2: Evaluation of lung function in the 5 weeks and 10 weeks models**. Mice are exposed either to room air (RA) and challenged with PBS, or exposed to cigarette smoke (CS) and challenged with poly(I:C), during 5 weeks (A-D) or 10 weeks (E-J). RA+PBS-exposed mice are represented by black circles and CS+poly(I:C)-exposed mice by gray squares. Data represent individual mice and are analyzed by Mann–Whitney test. (A) FEV0.05/FVC (FEV0.05: Forced Expiratory Volume during 0.05 s, FVC: Forced Vital Capacity). (B) FEV0.05. (C) FVC. (D) Peak Flow. (E) Inspiratory capacity (“IC”). (F) resistance of the respiratory system (“Rrs”). (G)

Compliance of the respiratory system (“Crs”). (H) Newtonian resistance (“Rn”). (I) Tissue damping (“G”). (J) Tissue elastance (“H”).

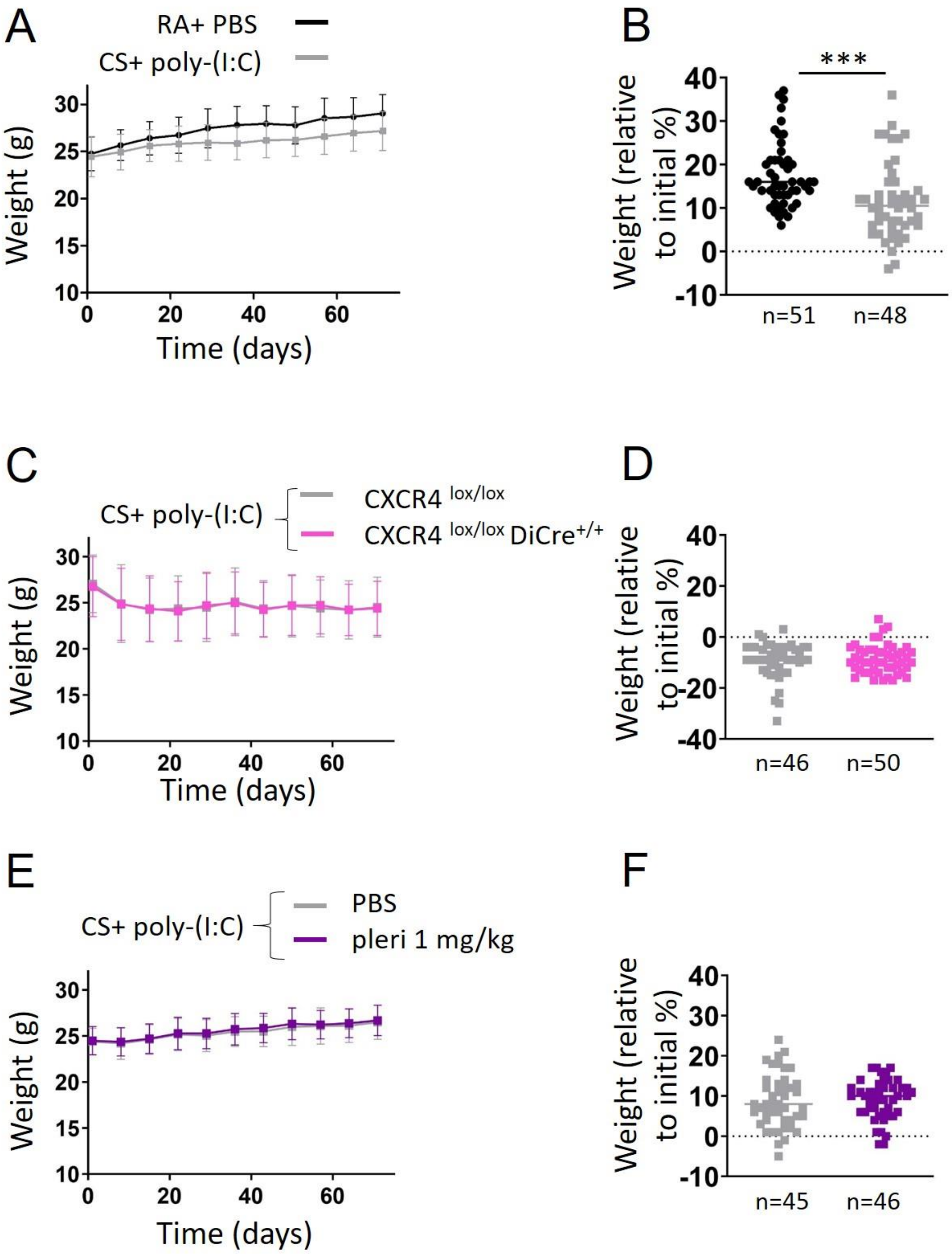


**Figure S3: Weight gain in the different protocols.** (A, C, E) Mean weight according to time during the protocol of experimental COPD (A), with conditional inactivation of CXCR4 (C) and with pharmacological inhibition of CXCR4 using plerixafor ("pleri", E). Error bars represent standard deviations. RA: Room air, CS: Cigarette smoke. (B, D, F) Weight gain, relatively to the initial weight, expressed in percentage during the protocol of experimental COPD (B), with conditional inactivation of CXCR4 (D) and with pharmacological inhibition of CXCR4 (F). Medians are represented as horizontal lines. ***: P<0.001. Unpaired t test or Mann-Whitney test.

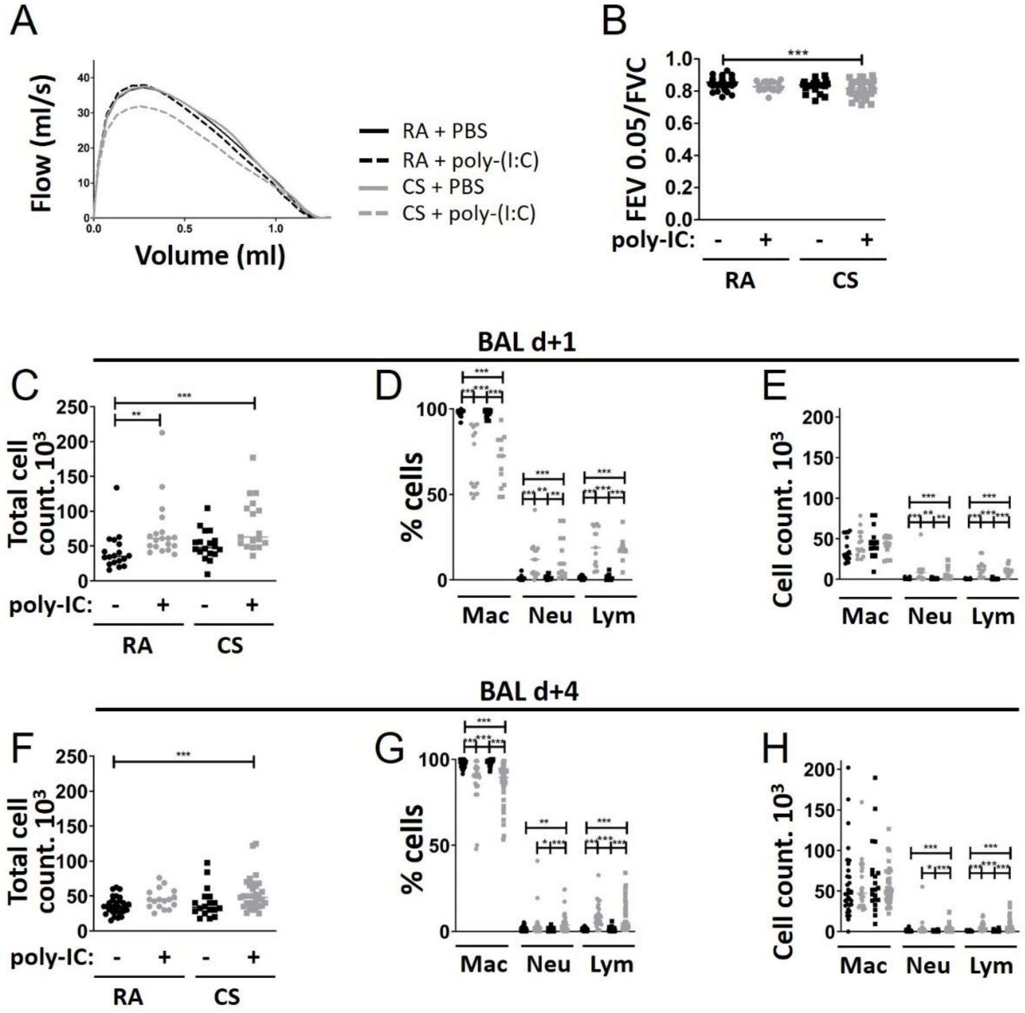


**Figure S4: Respective role of cigarette smoke exposure and poly(I:C) instillation in experimental COPD**. Mice are exposed either to room air (RA) and challenged with PBS (RA+PBS), exposed to RA and challenged with poly(I:C) (RA+ poly(I:C)), exposed to cigarette smoke (CS) and challenged with PBS (CS+PBS), or exposed to cigarette smoke (CS) and challenged with poly(I:C) (CS+poly(I:C)), during 10 weeks. They are sacrificed at day 68 or 71 (1 day and 4 days after the last poly(I:C) instillation). (A) Average expiratory flow-volume curves of mice in the different conditions. d) FEV0.05/FVC (FEV0.05: Forced Expiratory Volume during 0.05 s, FVC: Forced Vital Capacity). Data represent individual mice. ***: P<0.001. one-way analysis of variance followed by Bonferroni’s post-tests. RA+PBS: n=40, RA+ poly(I:C): n=17,

CS+PBS: n=19, CS+poly(I:C): n=41) (C, F) Total cell count in bronchoalveolar lavage (BAL) 1 day (C) and 4 days (F) after the last poly(I:C) instillation. (D-E, G-H) BAL differential cell recovery 1 day (D-E) and 4 days (G-H) after the last poly(I:C) instillation. (B, C-H) RA+PBS: black circles, RA+ poly(I:C): gray circles, CS+PBS: black squares, CS+ poly(I:C): gray squares. D+1: RA+PBS: n=18, RA+ poly(I:C): n=19, CS+PBS: n=18, CS+poly(I:C): n=17. D+4: RA+PBS: n=30, RA+ poly(I:C): n=17, CS+PBS: n=18, CS+poly(I:C): n=31. (C-H) *: $P<0.05$, **: $P<0.01$, ***: $P<0.001$. Kruskal-Wallis test followed by Dunn's post-tests.

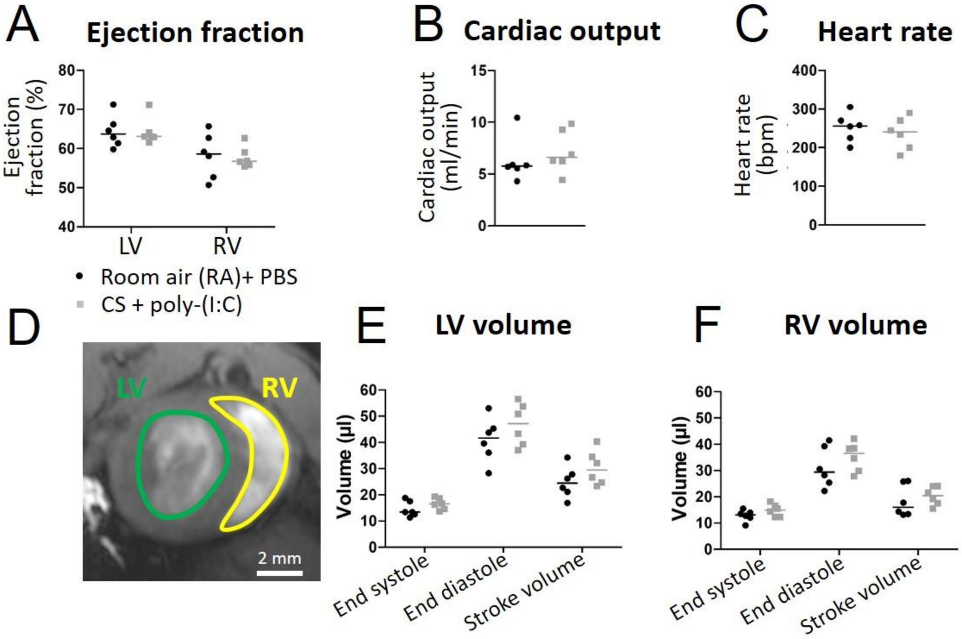


**Figure S5: Characterization of cardiac volumes in experimental COPD.** Mice are exposed either to room air (RA) and challenged with PBS (black circles), or exposed to cigarette smoke (CS) and challenged with poly(I:C) (gray squares), during 10 weeks. (A) RV and LV ejection fraction are similar between control and exposed mice (RV ejection fractions: respectively 64.3 and 64.3%, LV ejection fractions: respectively 58.2 and 58.8%). (B) Cardiac output. (C) Heart rate. (D) Representative short-axis cine magnetic resonance images at end-diastolic stage in control mice, showing the left ventricle (LV) and right ventricle (RV). (E) Volume of the left ventricle. (F) Volume of the right ventricle. Both are similar between control and exposed mice, indicating an absence of dilated cardiomyopathy. RA+PBS: n=6, CS+poly(I:C): n=6

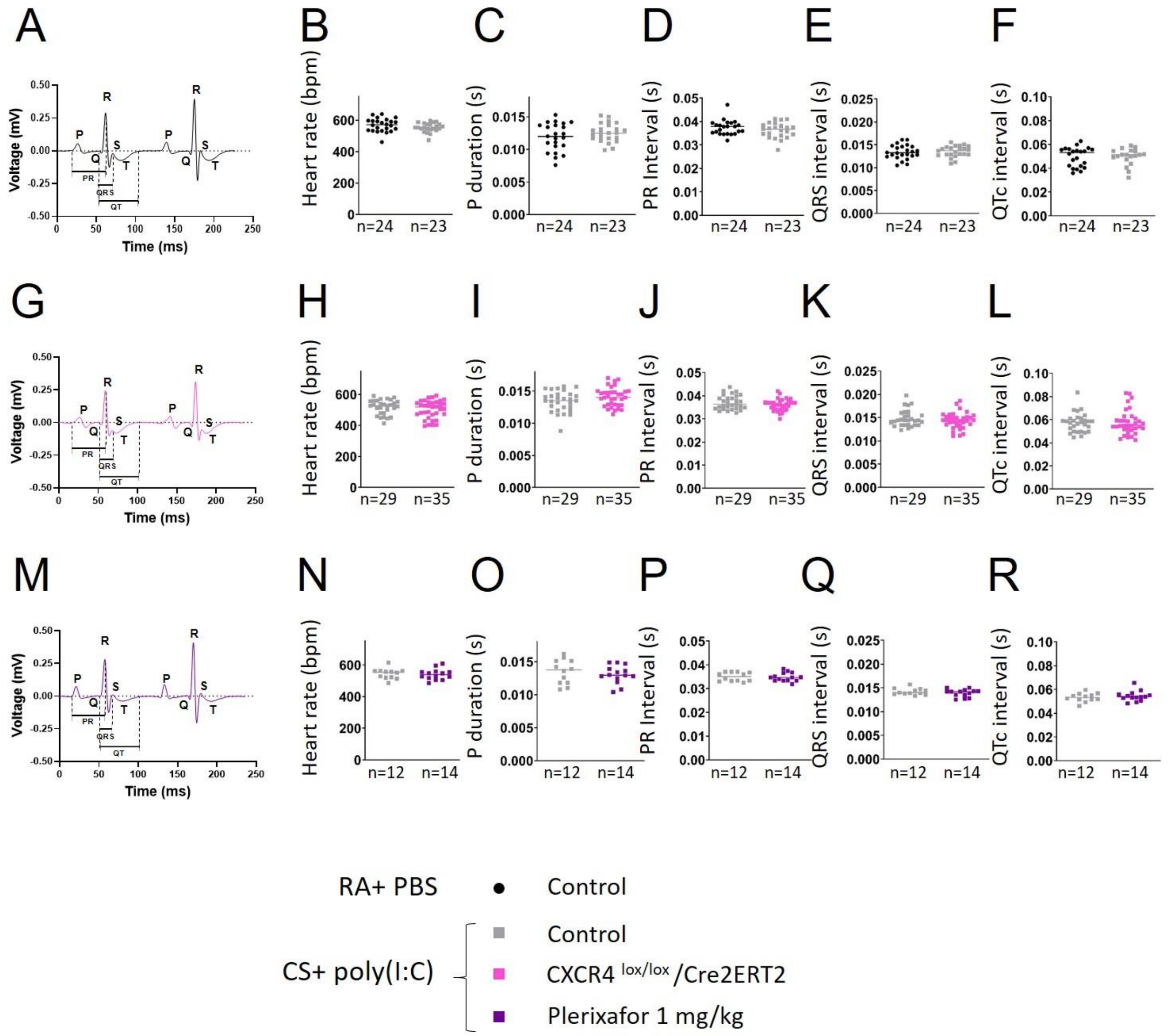


**Figure S6: Effect of CXCR4 inhibition on electrocardiogram (ECG) parameters in experimental COPD.** ECG parameters were quantified at the end of the protocol of experimental COPD (A-F), with conditional inactivation of CXCR4 (G-L) and with pharmacological inhibition of CXCR4 using plerixafor (M-R). RA: Room air, CS: Cigarette smoke. (A, G, M) Average ECG curves from one representative animal in the different conditions. (B, H, N) Heart rate. (C, I, O) Duration of P wave. (D, J, P) Interval between P wave and the beginning of the QRS complex. (E, K, Q) Interval of the QRS complex. (F, L, R) Corrected QT interval (QTc). The QTc was calculated using Mitchell formula (QT interval/(square root of the RR interval*10)). Medians are represented as horizontal lines. Unpaired t test or Mann-Whitney test.

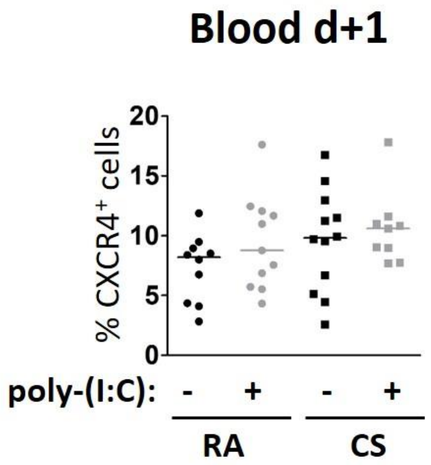


**Figure S7: Respective role of cigarette smoke exposure and poly(I:C) instillation on CXCR4 expression in circulating cells**. Mice are exposed either to room air (RA) and challenged with PBS, exposed to RA and challenged with poly(I:C), exposed to cigarette smoke (CS) and challenged with PBS, or exposed to cigarette smoke and challenged with poly(I:C) during 10 weeks. They are sacrificed at day 68 (1 day after the last poly(I:C) instillation). The graph respresents levels of circulating $CXCR4^{+}$ cells in the different conditions. RA+PBS: n=10, RA+ poly(I:C): n=11, CS+PBS: n=12, CS+poly(I:C): n=9. Data represent individual mice. Medians are represented as horizontal lines.

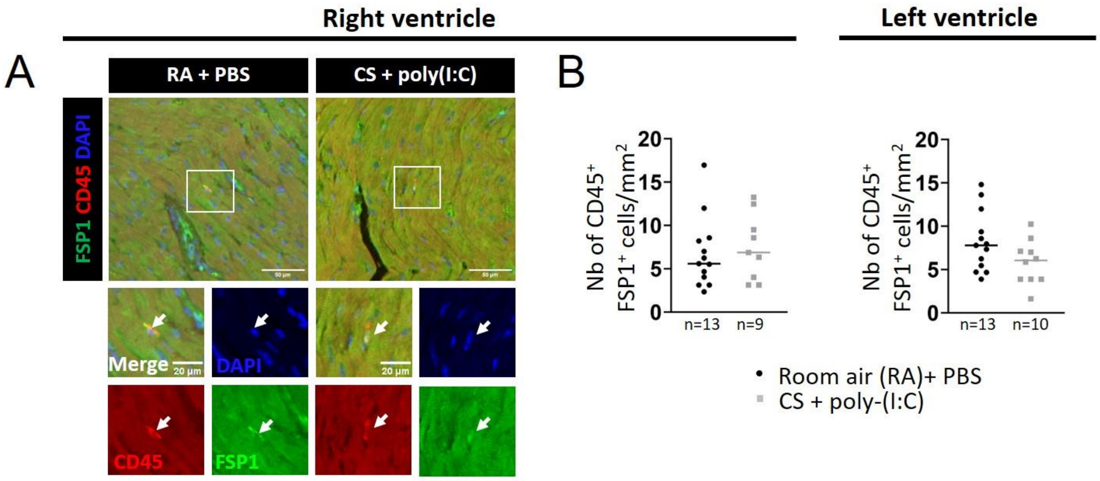


**Figure S8: Characterization of fibrocyte density in the heart in experimental COPD**. Mice are exposed either to room air (RA) and challenged with PBS, or exposed to cigarette smoke (CS) and challenged with poly(I:C), during 10 weeks. They are sacrificed at day 71 (i.e. 4 days after the last poly(I:C) instillation) (A) Representative stainings of CD45 (red) and FSP1 (green) in right ventricles of heart mice. The white arrows indicate fibrocytes, defined as $CD45^+$ $FSP1^+$ cells. (B) Quantification of fibrocyte density (number of fibrocytes normalized by total surface area). For the right ventricle, one sample was lost. Medians are represented as horizontal lines.

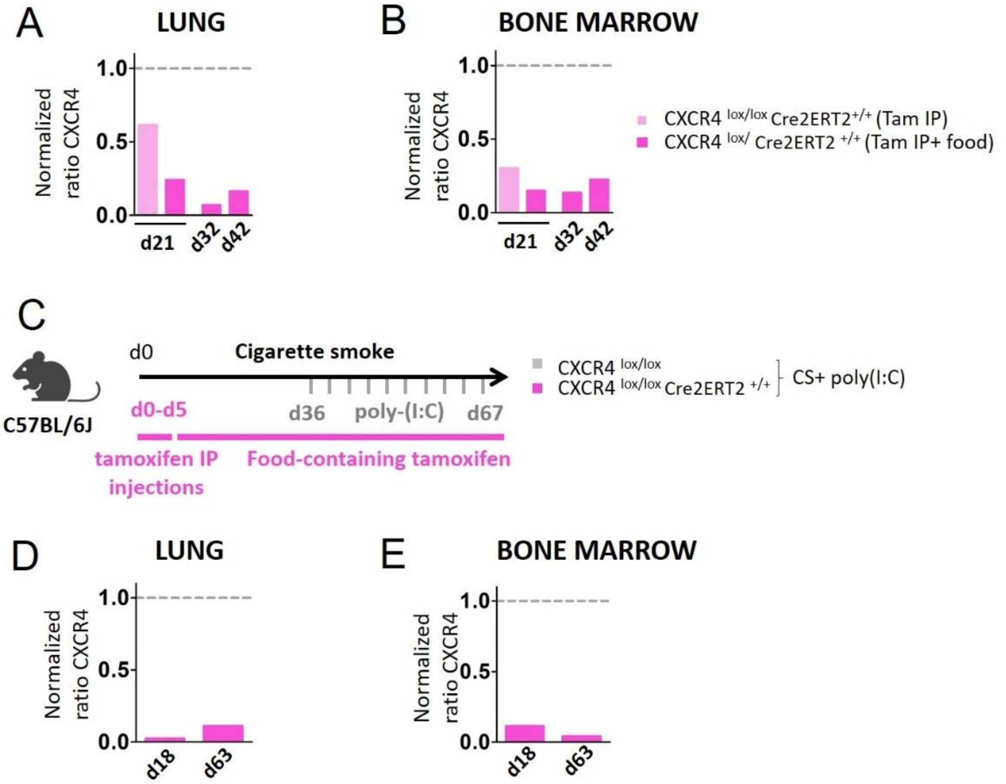


**Figure S9: Validation of CXCR4 conditional knockout in experimental COPD.** (A-B) $CXCR4^{lox/lox}$ mice and $CXCR4^{lox/lox}/Cre2ERT2^{+/+}$ mice were administered with 1 mg intraperitoneal tamoxifen for 5 consecutive days and then either fed with normal food (“Tam IP”, light pink, n=1), or with fed by tamoxifen-containing food (“Tam IP+food”, pink, n=1) and sacrificed at the indicated time points. CXCR4 mRNA levels measured by qPCR in lung (A) and bone marrow (B). The levels are normalized to those obtained in $CXCR4^{lox/lox}$ mice. (C) $CXCR4^{lox/lox}$ (n=1) and $CXCR4^{lox/lox}/Cre2ERT2^{+/+}$ mice (n=1) were administered with 1 mg intraperitoneal tamoxifen for 5 consecutive days at the beginning of the COPD protocol, and then fed by tamoxifen-containing food during the remaining protocol. They are sacrificed at day 18 or 63. (D-E) CXCR4 mRNA levels measured by qPCR in lung (D) and bone marrow (E). The levels are normalized to those obtained in $CXCR4^{lox/lox}$ mice.

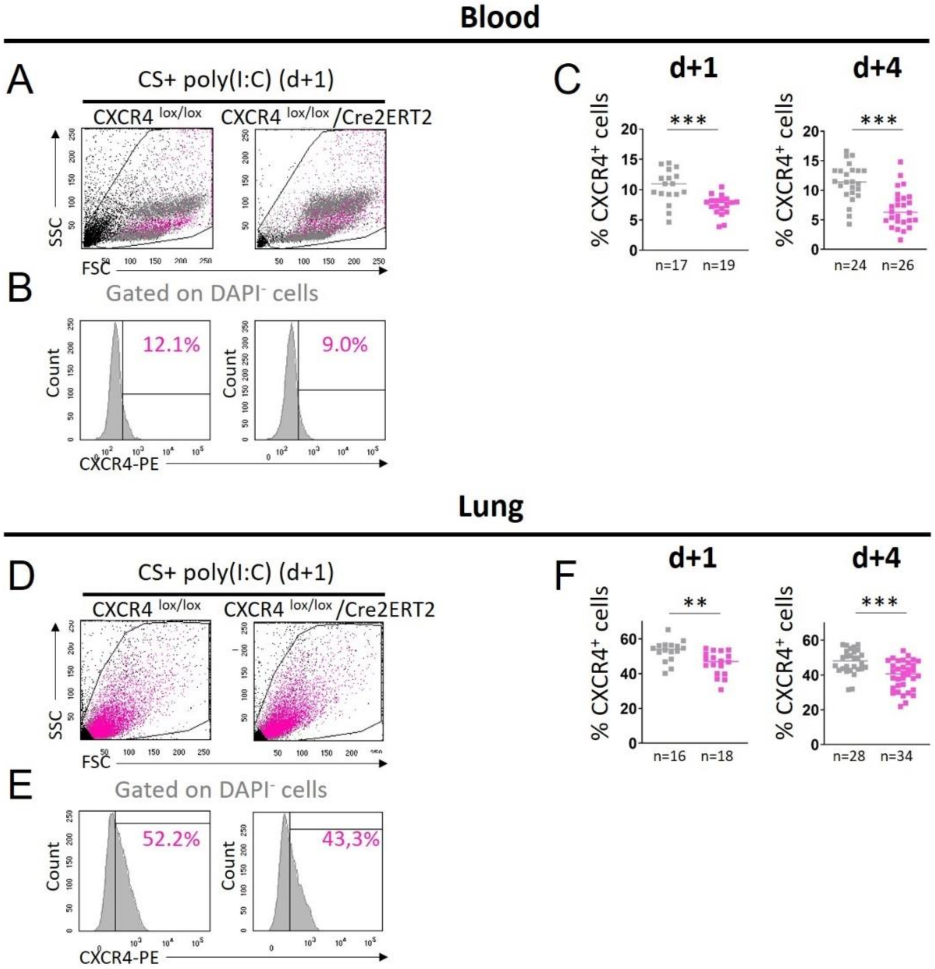


**Figure S10: Characterization of CXCR4 expression in the blood and lung upon conditional knockout of CXCR4 in experimental COPD.** CXCR4$^{lox/lox}$ (gray squares) and CXCR4$^{lox/lox}$/Cre2ERT2 (pink squares) mice are exposed to cigarette smoke (CS) and challenged with poly(I:C). All the mice were administered with 1 mg intraperitoneal tamoxifen for 5 consecutive days at the beginning of the COPD protocol, and then fed by tamoxifen-containing food during the remaining protocol. They are sacrificed at day 68 or 71 (1 day and 4 days after the last poly(I:C) instillation). (A, D) Dot plots represent representative side scatter (SSC, y-axis)-forward scatter (FSC, x-axis) graphs of circulating (A) and lung (D) cells. DAPI$^-$ cells and DAPI$^-$ CXCR4$^+$ cells are shown respectively in gray and pink. (B, E) Histograms represent representative cell count (y-axis) versus CXCR4-Phycoerythrin (PE) fluorescence (x-axis) in DAPI$^-$ circulating

(B) and lung (E) cells. Percentages of CXCR4$^+$ cells in the DAPI$^-$ cells are shown in pink. (C, F) Levels of circulating (C) and lung (F) CXCR4$^+$ cells 1 day (d+1) and 4 days (d+4) after the last poly(I:C) instillation. (C, H) Medians are represented as horizontal lines. **: P<0.01, ***: P<0.001. Unpaired t test or Mann-Whitney test.

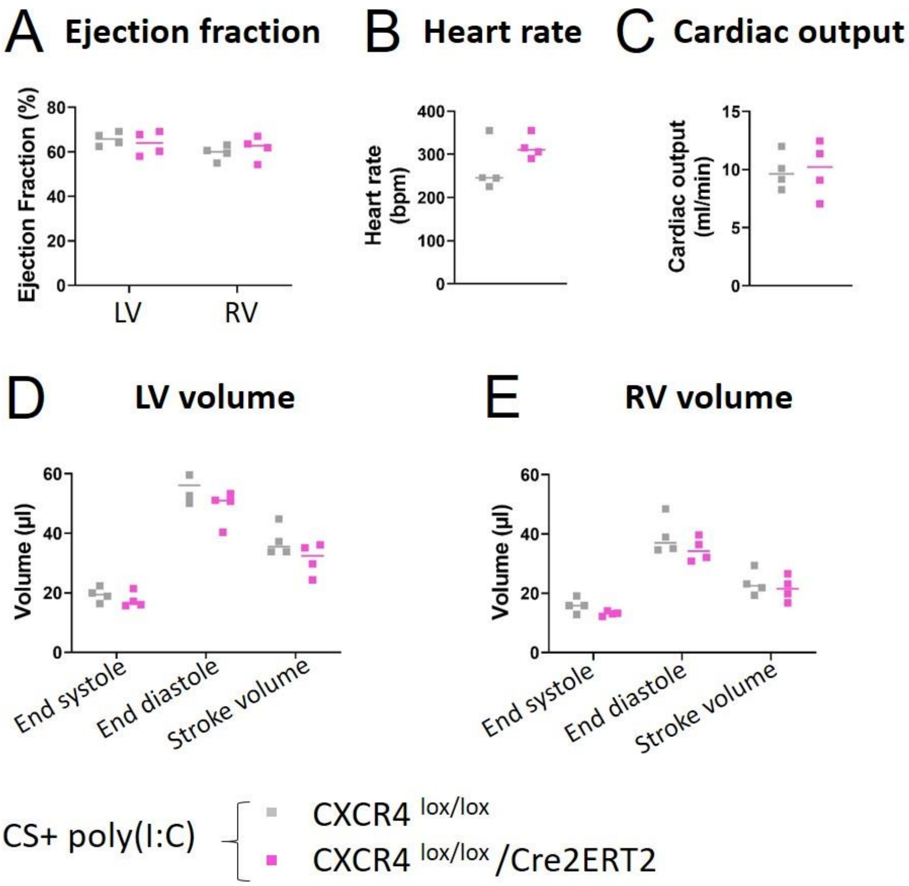


**Figure S11: Characterization of cardiac function and anatomy by MRI upon conditional knockout of CXCR4 in experimental COPD.** CXCR4$^{lox/lox}$ mice (gray squares, n=4) and CXCR4$^{lox/lox}$/Cre2ERT2 mice (pink squares, n=4) are exposed to cigarette smoke (CS) and challenged with poly(I:C). (A) Ejection fraction. (B) Heart rate. (C) Cardiac output. (D) Volume of the left ventricle (LV). (E) Volume of the right ventricle (RV). Mann-Whitney test.

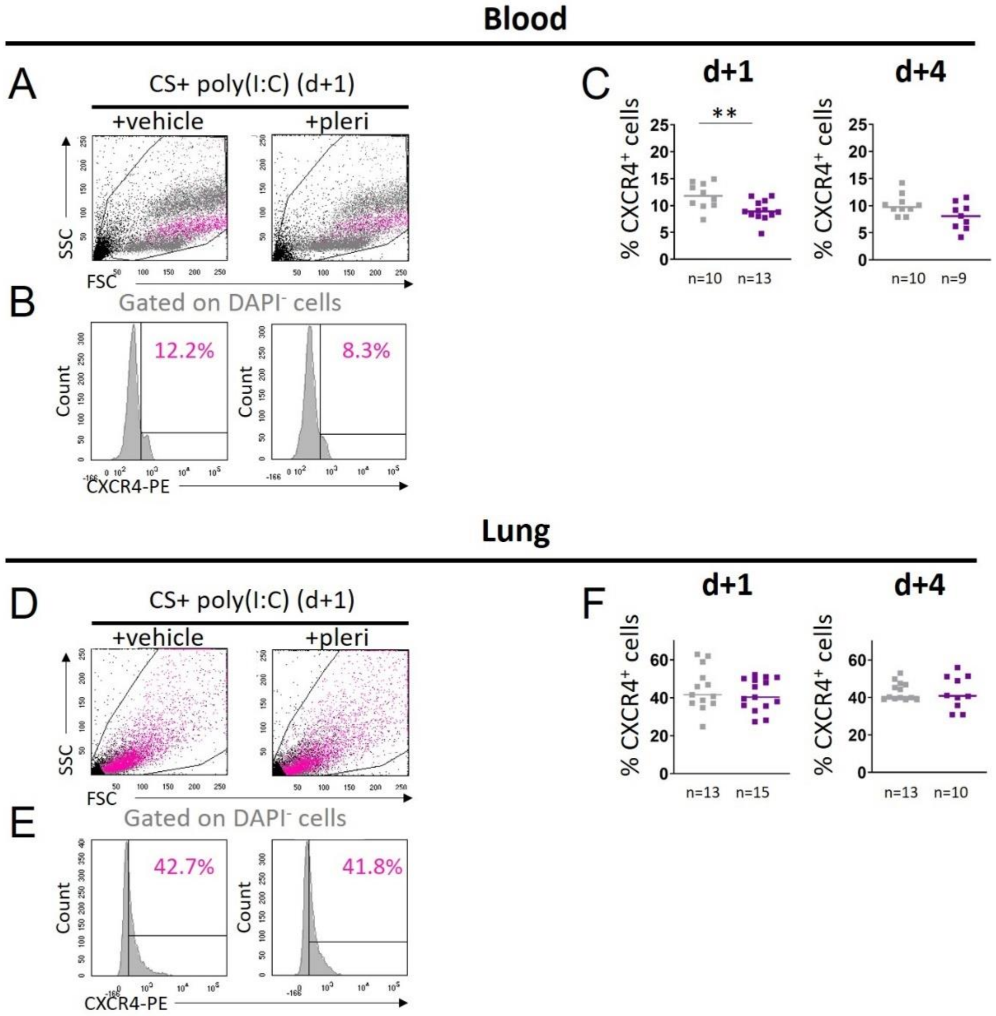


**Figure S12: Characterization of CXCR4 expression in the blood and lung upon pharmacological blockage of CXCR4 in experimental COPD.** Mice are exposed to cigarette smoke (CS), challenged with poly(I:C) and treated with plerixafor ("pleri", 1 mg/kg, purple squares) or its vehicle (gray squares). They are sacrificed at day 68 or 71 (1 day and 4 days after the last poly(I:C) instillation). (A, D) Dot plots represent representative side scatter (SSC, y-axis)-forward scatter (FSC, x-axis) graphs of circulating (A) and lung (D) cells. DAPI$^{-}$ cells and DAPI$^{-}$ CXCR4$^{+}$ cells are shown respectively in gray and pink. (B, E) Histograms represent representative cell count (y-axis) versus CXCR4-Phycoerythrin (PE) fluorescence (x-axis) in DAPI$^{-}$ circulating (B) and lung (E) cells. Percentages of CXCR4$^{+}$ cells in the DAPI$^{-}$ cells are shown in pink. (C, F)

Levels of circulating (C) and lung (F) CXCR4$^+$ cells 1 day (d+1) and 4 days (d+4) after the last poly(I:C) instillation. (C, F) Medians are represented as horizontal lines. **: $P<0.01$. Unpaired t test or Mann-Whitney test.

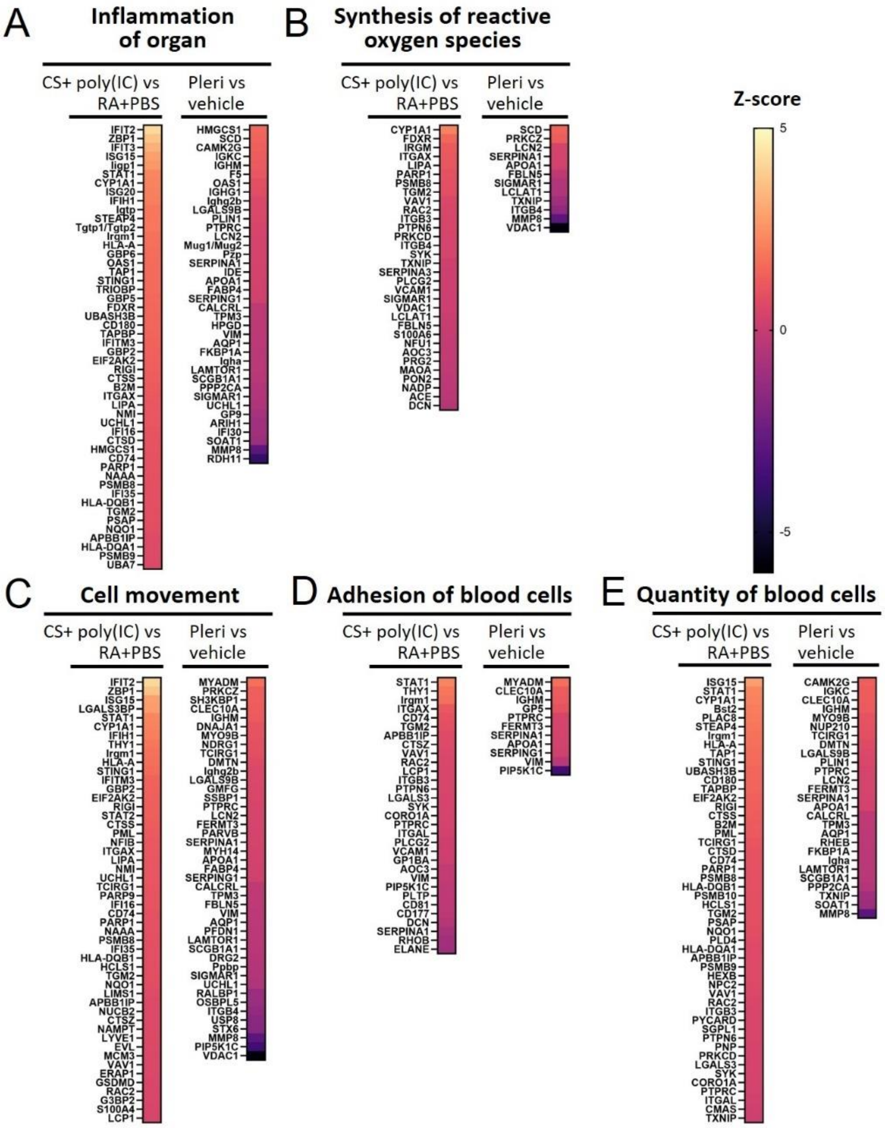


**Figure S13: Comparison analysis of proteome changes in experimental COPD and upon pharmacological blockage of CXCR4.** Heatmaps of differentially regulated proteins in CS+poly(I:C) (n=5) *vs* RA+PBS (n=5)-exposed lungs (n=5) (all treated by vehicle) and plerixafor (n=5) vs vehicle (n=5)-treated lungs (all exposed to CS+poly(I:C)), from the pathways “Inflammation of organ” (A), “Synthesis of reactive oxygen species” (B), “Cell movement” (C), “Adhesion of blood cells (D), “Quantity of blood cells (E). The color scale indicates the $\log_2$ fold changes of abundance for each protein.

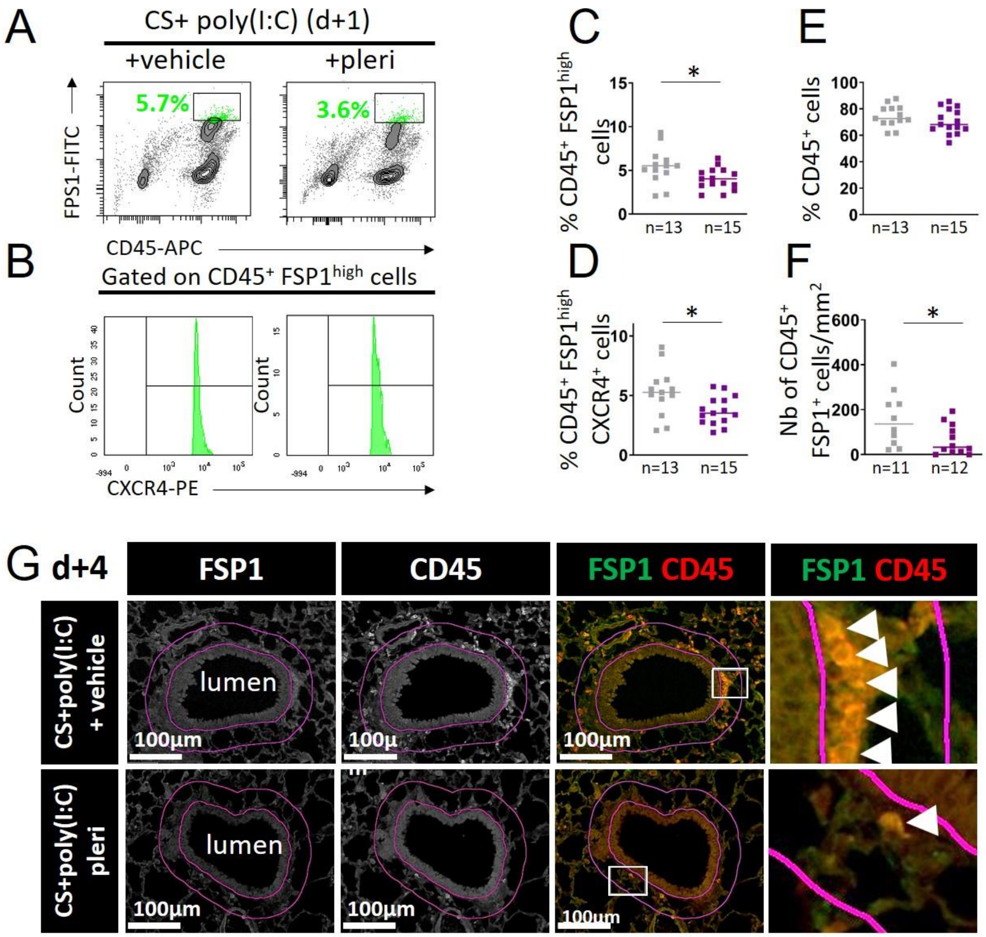


**Figure S14: Characterization of fibrocyte level and peri-bronchial density in lung upon pharmacological blockage of CXCR4 in experimental COPD**. Mice are exposed to cigarette smoke (CS), challenged with poly(I:C) and treated with plerixafor ("pleri", 1 mg/kg, purple squares) or its vehicle (gray squares). They are sacrificed at day 68 or 71 (1 day and 4 days after the last poly(I:C) instillation). (A) Representative flow cytometry contour plots in each condition. The x-axis is for CD45-Allophycocyanin (APC) fluorescence and the y-axis, FSP1-Fluorescein-5-isothiocyanate (FITC) fluorescence. Percentages of $CD45^+$ $FSP1^{high}$ cells are shown in green. (B) Representative cell count (y-axis) versus CXCR4-Phycoerythrin (PE) fluorescence (x-axis) in $CD45^+$ $FSP1^{high}$-cell subsets in mouse lungs. (C-E) Levels of lung $CD45^+$ $FSP1^{high}$ cells (c), $CD45^+$

$FSP1^{high}$ $CXCR4^+$ cells (D), $CD45^+$ cells (E) 1 day after the last poly(I:C) instillation in each condition. (F) Quantification of fibrocyte density (normalized by peri-bronchial area) 4 days after the last poly(I:C) instillation in each condition. (G) Representative stainings of CD45 (red) and FSP1 (green) in peri-bronchial area (delimited in pink) in each condition. The white arrows indicate fibrocytes, defined as $CD45^+$ $FSP1^+$ cells. (C-F) Medians are represented as horizontal lines. *: $P < 0.05$, Unpaired t test or Mann-Whitney test.
